# Attention-based frontal-posterior coupling for visual consciousness in the human brain

**DOI:** 10.1101/2024.06.30.601105

**Authors:** Xilei Zhang, Chao Zhang, Xiqian Wu, Wenjing Zhou, Sheng He, Yi Jiang, Kai Zhang, Liang Wang

## Abstract

We usually perceive what we are attending to. How goal-directed attention contributes to conscious perception remains yet elusive. Here we combined a novel psychophysical paradigm with intracranial electroencephalography data to investigate this issue in the human brain. Relative to unattended conditions, goal-directed attention modulated early activity and inter-regional connectivity, even though this part of attention failed to predict image detectability. Later, the coupling between the frontal and posterior brain got established and maintained but the signals exchanged did not inform fine-grained image contents but instead reflect success or failure of attentional capture. This part of captured attention proportionally predicted image detectability. These results attribute consciousness to attention-based coupling between the frontal and posterior brain as a whole, rather than activity of either part alone.

## Main text

The emergence of subjective consciousness in the brain presents a multifaceted puzzle, with attention occupying a central position. Despite the daily experience that what we consciously perceive are usually what we are attending to, ongoing debate revolves around the question of whether some interaction between attention and sensory processing is necessary for conscious perception, and if so, which type of attention? (*1–4*). In terms of neural mechanisms, though goal-directed attention dramatically changes both pre-stimulus and stimulus-evoked neural activity and connectivity(*5–16*), the precise processes through which this goal-directed attention finally facilitates (if not initiates) conscious perception still remain elusive.

At the first step, we intuitively proposed it may be the part of attention intertwining with specific sensory processing that correlates with visual consciousness of this stimulus, rather than the part of attention generally directed to stimulus location beforehand. To test this possibility, via a psychophysical experiment, we newly designed a paradigm separately quantifying how much attention has been pre-allocated to the image location before stimulus onset, and how much attention has been further captured by the image when it is consciously seen. As revealed using this paradigm, goal-directed attention is indeed a prerequisite to consciously differentiate the visible versus invisible images. However, it is the captured attention, which intrinsically reflects the interaction between attention and visual processing, that correlates with image detectability (i.e., a measure of subjective visual experience). Previously, attentional capture was usually regarded as a mechanism competing with goal-directed attention (*17–20*). Here, the present study provides new evidence on their joint collaboration in supporting visual consciousness.

Then, combining this paradigm with high spatial-temporal resolution iEEG (*21*) data, we further searched for and identified attention-induced neural dynamics which reflect the effect of goal-directed attention and captured attention, respectively. Especially, we found the neural marker of attentional capture process appeared till to roughly 300ms post-stimulus onset in the attended condition only. This identification thus allowed us to discriminate early neural dynamics tied to goal-directed attention from late neural dynamics tied to captured attention, so as to evaluate what neural dynamics make necessary preparation for later processes but do not directly correlate with consciousness, and what more likely reflect how consciousness arises in the human brain. Currently, fierce debates (e.g., (*22*)) are arguing to attribute consciousness to either the frontal brain (*23–25*) or the posterior brain (*26–30*). Neural findings about the late-stage neural dynamics would also give these consciousness-related theories an empirical test.

### Experiment 1: Joint collaboration between goal-directed attention and captured attention in behavior

We designed a highly demanding counting task (**Fig.1A, 1B**, Method) in which the performance would change as a function of the amount of consumed attention. During counting, an image was suddenly presented in the peripheral visual field. Since the amount of attentional resources is limited (*31*), if attention is somehow distracted by an image, less attention available for the counting task would lead to a deteriorated counting performance.

**Figure 1.**
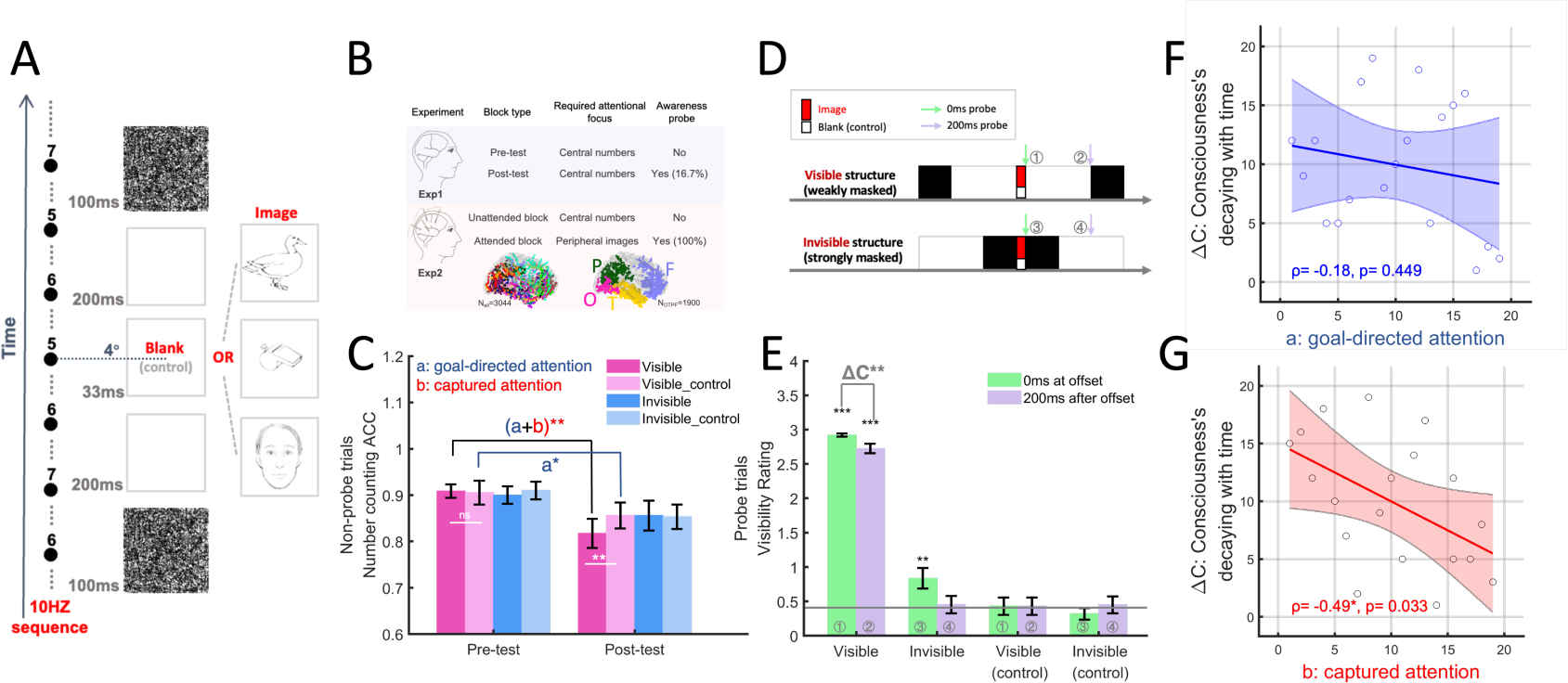
Captured attention correlates with visual consciousness. (A) Schematic illustration of the timing structure of an exemplar visible trial consisting of a central number stream (10 Hz) and a peripheral image stream, displayed in parallel. (B) The supply of attention to the peripheral image was manipulated across two independent experiments. Intracranial local field potential signals were only recorded in experiment 2, of which regions of interest consisted of four neocortex lobes (including 1900 implanted contacts marked in different color). (C) Attention deliberately directed to peripheral image (component a) and attention passively captured by visible images (component b) were indirectly measured by number counting performance. (D) In 16% randomly selected trials (post-test of Exp.1), an awareness probe was introduced at either 0ms or 200ms post-stimulus offset, in order to spare partial attention for peripheral image. Black and white squares indicate visual mask and blank screen, respectively. (E) Visibility rating for these probes revealed how much subjective visibility decays over 200ms interval. Spearman correlation analysis revealed a negative correlation between this decay and attention for captured attention (G), but not for goal-directed attention (F), suggesting captured attention plays critical roles in visual consciousness. See **Fig.S2** for replication study of this effect in an independent sample.

In the pretest session (i.e., without probe, **movie S1**), we required participants to focus on counting task and neglect the peripheral image completely. The image was either invisible (heavily masked) or visible (weakly masked). As a result, counting performance did not vary with visibility (Fig. 1C, F(_3, 54_)= 0.16, p=0.922), suggesting that in the pretest session peripheral salient images failed to capture attention.

In the posttest session (i.e., with probe**, movie S2**), we still required participants to set priority the counting task and perform at their best. The only difference was that a probe testing for visibility of the peripheral image was occasionally introduced in 16% randomly selected probe trials (**Fig.1D**, Method). In all trials (including the rest 84% non-probe trials), we anticipated participants would in fact spare attention according to two goals. Though the primary goal was counting, we assumed this manipulation would induce a secondary goal. That is, participants would shift some attention away from the counting task, allocating partial attention to peripheral image intentionally (hereafter we call this part of attention as ‘goal-directed attention’ for the image), to cope with occasional probe test. If so, counting performance of non-probe trials (see following) would decrease because the attention resource available for the counting task was reduced (Method).

In 84% non-probe trials, there was indeed a general decrease (**Fig.1C**, t(_18_)=-3.57, p=0.002) on counting performance in the posttest session relative to the pretest baseline. This decrease was not an artifact of fatigue since, in each session, task performance did not decrease as a function of sequential order of blocks in both the pretest (2 blocks, t(_18_)=-1.45, p=0.164) and posttest sessions (10 blocks, F(_9,162_)=1.16, p=0.322). Notably, visible image trials relative to invisible image trials encountered an additional decrease (interaction: F(_1, 18_)=6.82, p=0.018) which was irrespective of categorical contents (F(_2,36_)=0.78, p=0.464). Probe trials were not counted into this calculation so this part of attention was not captured by the probe cue. By contrast, an invisible signal failed to capture attention because performance of the invisible image trials was equivalent with the other two control conditions where only masks were displayed in the absence of the target image. This finding indicated that when attention became available, conscious signal (visible versus invisible) automatically captured attention. To avoid any confounding from masks, we subtracted the amount of attention generally divided to visible control trials (component **a:** goal-directed attention, t(_18_)=2.65, p=0.016) from that of visible image trials (component **a+b:** goal-directed plus captured attention, t(_18_)=3.62, p=0.002), obtaining a clean measure of the amount of captured attention (component **b**). Next, this measure was used to evaluate how the attention captured by the visible image contributed to its visual detectability.

Among 16% probe trials, a probe cue appeared either immediately at 0ms or shortly later at 200ms after image disappeared (**Fig.1D**). Whenever seeing the probe cue, participants orally reported whether the image displayed at that moment was ‘seen’ and if yes, they needed to report details about its category and identity (recorded online, **Fig.S1** and Methods). Above-chance category classification accuracy affirmed that visible and invisible images were indeed consciously and unconsciously perceived, respectively (**Fig.S1B**), while the decrease of seen ratio (**Fig.1E**, t(_18_)=-3.02, p=0.007) for 200ms versus 0ms probes indicated that the subjective visibility was decaying over a 200ms temporal gap. Calculated using probe and non-probe trials respectively, the magnitude of this decaying over time was negatively correlated with the amount of captured attention (spearman ρ= −0.49, p=0.033, **Fig. 1G**) but non-significant for goal-directed attention (**Fig.1F**, spearman ρ= −0.18, p=0.449). This finding had been well replicated (**Fig.S2**) in another experiment performed using the same paradigm and comparable sample size but recruiting different experimenter and audio encoders. These findings highlight a joint collaboration between goal-directed attention and captured attention in our paradigm. Once partial goal-directed attention is deliberately directed to stimulus location before stimulus onset, then visible image later becomes able to capture attention and this captured attention proportionally contributes to conscious visual perception of the image.

### Experiment 2: Joint collaboration between goal-directed attention and captured attention in the human brain

Experiment 2 aimed to further unlock how goal-directed attention and captured attention play collaborative roles during visual consciousness in the human neocortex by contrasting an unattended condition (number counting task, **movie S3**) versus a fully attended (category classification task, **movie S4**) condition. We recruited 21 drug-resistant epilepsy patients (**Fig.1B** lower panel, **Table S1**), who were implanted a number of iEEG electrodes (3,044 contacts in total) solely based on clinical demands.

### Spatial-temporal activation and functional disconnection between the frontal and posterior brain at the unattended condition

We first looked at neural responses at the unattended condition when peripheral images were in a case of inattentional blindness (*32, 33*). Replicating behavioral results of Exp1, visible signals failed to capture attention relative to the invisible signals in behavior (**Fig.S3**). In terms of neural response, taking a representative contact in the primary visual area (V1) as an example (**Fig. 2A**, responsive for contralateral image; **Fig.S4**, non-responsive for ipsilateral image), V1 response to invisible images was weak and transient (amplitude 40µV, duration 10ms, upper panel) while response for visible images (400µV, 460ms) was about ten times stronger and much longer (**Fig.2A**, middle and bottom panels). This finding provides direct evidence favoring the existence of recurrent processing despite lack of attention (*34*).

**Figure 2.**
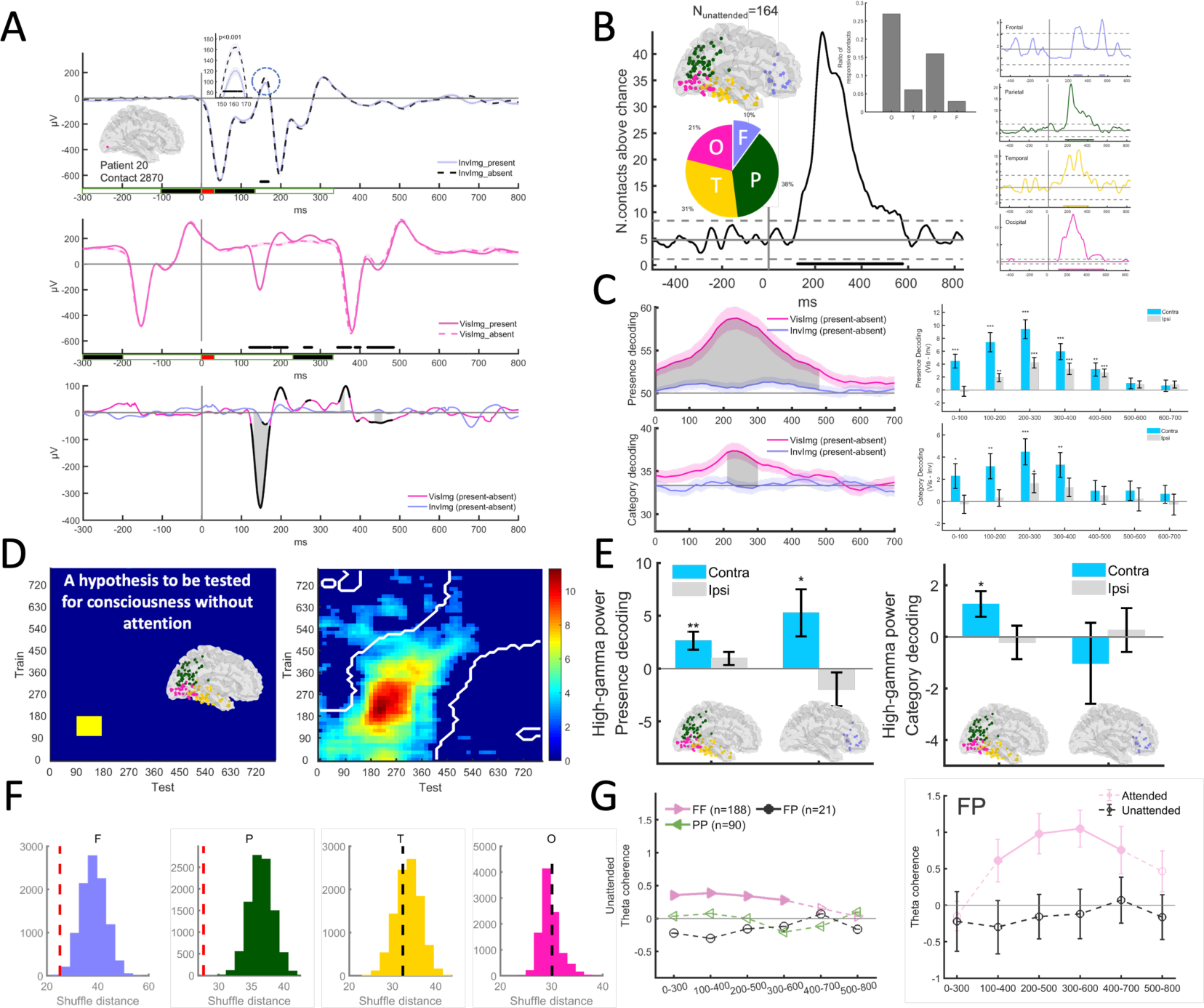
Preconscious visual processing in the unattended condition. (A) An exemplar responsive contact (MNI=[2.1, −88.5, −5.5]) at the primary visual cortex (V1) showed a weak and transient response to an invisible image, whereas a visible image evoked a much stronger and longer response even though attention was directed elsewhere. At the bottom of the x-axis, rectangles face-colored in white, black and red indicate the presentation of a blank screen, a visual mask and a target image (or a blank control), respectively. (B) Responsive contacts (N=164) extended to widespread regions of the posterior brain and a small subset of the frontal brain. Gray dotted lines indicate threshold of chance level (see Methods). For each lobe of interest, the pie and bar charts show the percentage among all responsive and implanted contacts, respectively. The rightmost panel shows the number of responsive contacts for each lobe. (C) Posterior activity (pooling responsive contacts at O, T and P lobes) decoded both coarse-grained binary presence and fine-grained categorical contents of the consciousness-related signal (left panel), especially by the contralateral hemisphere (right panel). (D) Coarse-grained representations generalized homogeneously across time for approximately 450 ms (right panel), opposing the hypothesis (left panel) for transient existence of consciousness without attention and suggesting a preconscious state between unconscious and conscious. (E) A small subset of the frontal brain was also responsive in the preconscious state, with its high-gamma activity decoding coarse-grained representations only, unlike the posterior brain which decoded both coarse-grained and fine-grained representations. (F) In the frontal and parietal lobes, responsive contacts were closely clustered with a significantly smaller mutual distance than shuffled chance. (G) Frontal-parietal (FP) theta coupling was not established in the unattended condition (left panel); on the contrary, FP theta coupling got established only when attention was engaged (right panel).

Beyond V1, RP activity in the posterior brain also expanded to broader regions of the occipital (O), temporal (T) and parietal (P) lobes (**Fig.2B, Fig.S5**, **movie S5**). This posterior activity encoded both binary presence (**Fig.2C**, upper panel) and categorical contents (**Fig.2C**, lower panel) of consciousness-related (visible versus invisible) signal, especially by slow theta oscillations (**Fig.S6**). Importantly, opposing the hypothesis for transient existence of consciousness without attention (**Fig.2D**, left panel), neural representations of binary presence were temporally homogenous across a duration as long as up to 450ms (**Fig.2D**, right panel). This length far exceeds ‘a conscious but transient and non-reportable state (*30, 34*))’. As experienced in the demo video (**movie S1**), when you are fully concentrating on counting from the fast-alternating number sequence you cannot explicitly ‘see’ a blurred peripheral image presented durably for 450ms (unless you pay attention to the image). Alternatively, findings above preferentially favor the tripartite distinction viewpoint (*34*) (dating back to Freud (*35*)) that salient stimuli enter into a ‘preconscious’ buffer state inter-between unconscious and conscious if not accessed by attention, i.e., ‘visible yet not seen’ (*34*).

A small set of the frontal brain was also activated in the preconscious state (**Fig.2B, Fig.S5**, **movie S5**), encoding the presence (200-400ms, **Fig.2E** left panel) rather than the contents (right panel) of the preconscious signal by the power of high-gamma oscillations (70-150 HZ), whose activity was potentially associated with local neuronal spiking (*36, 37*). Furthermore, these frontal responsive contacts clustered with a significantly smaller mutual distance than chance level (**Fig.2F**, permutation test, p=0.0042). This spatial clustering effect was also observed for parietal responsive contacts (**Fig.2F**, permutation test, p<0.0001, Table.S2) but not for occipital and temporal ones. Simultaneously, the peripheral salient signal failed to establish functional coupling (in theta band peaking at 4Hz, **Fig.S7**) between the frontal and posterior brain (**Fig.2G**, right panel showing frontal-parietal disconnection in the unattended condition). Given FP coupling is usually assumed to be associated with conscious perception (*34*), their disconnection further implies the image was processed in a preconscious buffer state. In this state, the peripheral salient image might have produced an ‘attend-to-me’ signal, reaching functionally specialized regions of the frontal and parietal lobes (particularly the frontal-parietal network and dorsal attention network, **Fig.S8**) to request attention (*19*), but this request had just been overridden to prevent actual visual distraction ((*18–20*), **Fig.1C** and **Fig.S3A**).

### Effect of goal-directed attention on early activity and connectivity before frontal-posterior coupling

In the attended condition, the number of responsive contacts sharply increased relative to the unattended condition (**Fig.3A**), with the frontal brain, a critical hub controlling attention (*6–8*), showing the greatest increase (**Fig.3B**, **movie S5**). Pre-stimulus alpha band power of the posterior brain was lateralized to the hemisphere ipsilateral to the attended side (**Fig.S9**), indicating that attention had been pre-allocated to the peripheral image location as required (*38*). Also, expectedly, this pre-allocated attention rendered the posterior brain into an elevated baseline state, boosting post-stimulus early visual activity even within the first 100ms (for review see (*5, 39*), **Fig. 3C** replicating the early effect at the posterior brain, **Fig.3D** showing similar effect at the frontal brain but of an about 100ms time lag).

**Figure 3.**
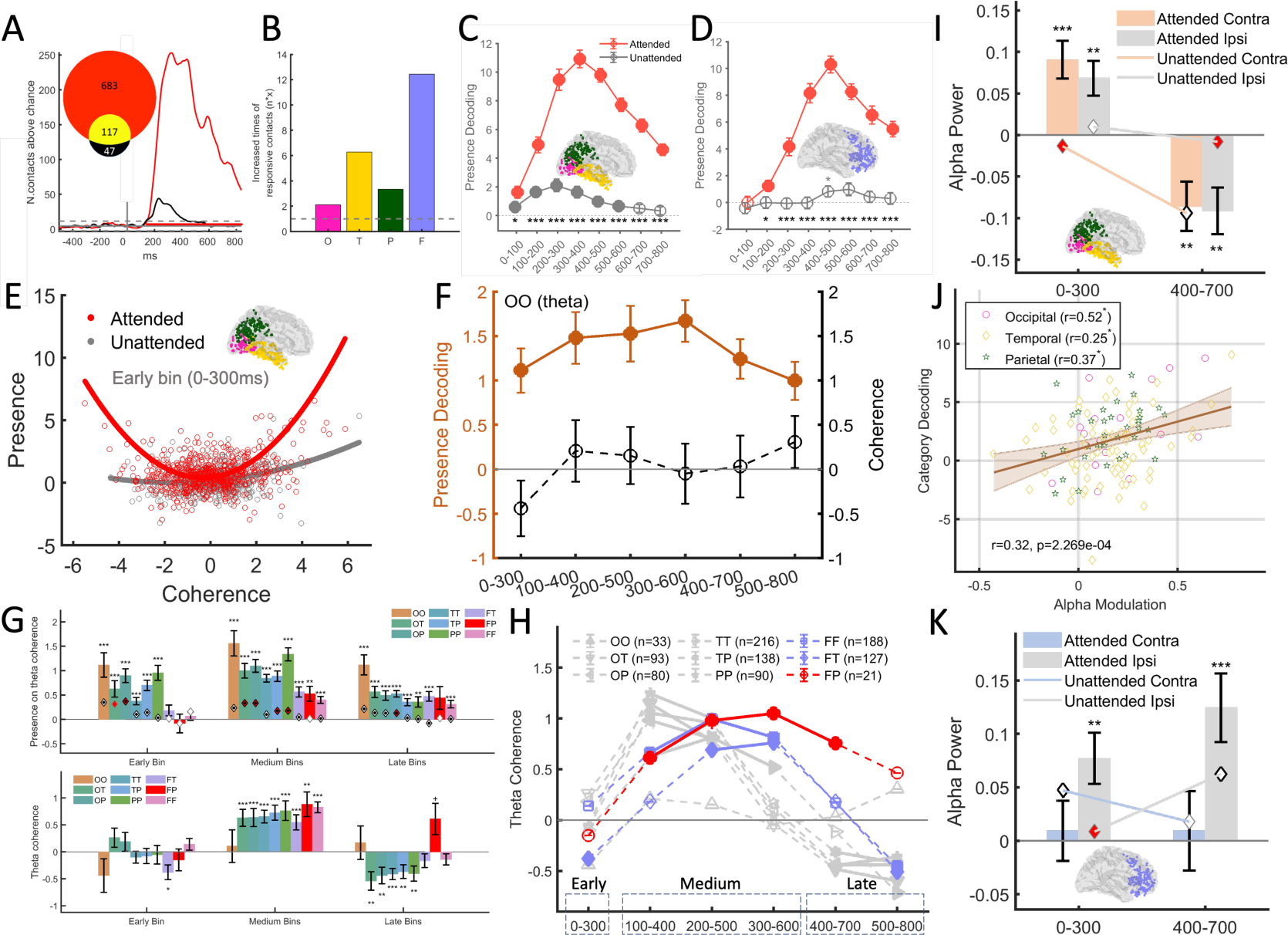
Neural dynamics switching early attentional modulation to late attentional capture. (A) Due to attentional engagement (the red line), the number of responsive contacts significantly increased relative to the unattended condition (the black line), with (B) the frontal brain showing the largest increase. Decoding performance (±SEM) of binary presence of consciousness was enhanced within the first 100 ms in the posterior brain (C), followed by a similar enhancement in the frontal brain about 100 ms later (D). (E) In terms of functional connectivity (calculated as adjusted imaginary coherence which may produce values exceeding 1, see Method), theta-phase coupling (positive values on the x-axis) and decoupling (negative values) coexisted intra-between the posterior brain in the early bin (0-300 ms), cancelling out each other to zero coherence by average. (F) Such functional reorganization effect was particularly observed for OO pairs which persisted till to the late stage. Coherence values and coherence-based presence decoding performance were tied to the right and left axis, respectively. (G) Despite these early changes intra-between the posterior brain, neither coupling nor decoupling occurred between the posterior and frontal brain at the early bin (±SEM). Subsequently, all four lobes exhibited coupling at medium bins, followed by decoupling at the late bins. Diamond symbols (as marked in **Fig.S15** and **S17**) indicate results in the unattended condition. (H) FP pairs maintained positive coupling for the longest duration, persisting into the late bin when other pairs became decoupled. (I) Throughout the whole-time course, attention induced a two-stage reversed alpha power pattern in the posterior brain (±SEM), observed in both hemispheres. (J) For contacts pooled from all three posterior lobes (brown fitting line) or each individual one (legend box), the amplitude of this reversed modulation effect positively correlated with the capacity in decoding categorical consciousness contents. *, p<0.05. (K) Conversely, in the frontal brain, alpha power (±SEM) was continuously lateralized to the hemisphere ipsilateral to the peripheral image side.

Beyond that, intra-between the posterior brain and at 4Hz peaked theta band (**Fig.S7**), we found attention had induced both coupled and decoupled pairs simultaneously at an early bin (0-300ms post-stimulus onset), with each sensitively detecting the salient emerging signal but leading to an overall zero coherence since they cancelled out each other (**Fig.3E**, red and gray curves for the attended and unattended condition, respectively; **Fig.S10** rigorously testing this U-shape relationship). For OO pairs (**Fig.3F**), notably, coexistence of decoupling and coupling even persisted across time till to late stages. Prior studies revealed similar attention-induced decoupling at the neuronal spiking level (*9, 10*). Extending this mechanism to a macroscopic scale in the human brain, the findings here suggest the goal-directed attention starts to exert its impact at the posterior brain by disrupting old connectivity patterns and creating new ones, especially starting from the early time.

Despite these early altered posterior activities and connectivity, we found the posterior brain was, however, still functionally isolated from the frontal brain at the same early bin. Frontal-posterior coherence (including the FP pair) neither increased nor decreased in response to the emerging signal (**Fig.3G**), akin to their isolation in the unattended condition (**Fig.2G**), suggesting consciousness had not yet emerged and the image (albeit attended) were still early processed in a preconscious state. In Exp.1, we have shown that attention supplied does not equal to attention captured and it is the captured attention that correlates with visual detectability (**Fig.1C, 1F** and **1G, for replication study see Fig.S2**). Findings here further show that these two types of attention operate differently in time. At the earliest stage, though the goal-directed attention has induced fundamental neural change in both the frontal brain and the posterior brain, attention has not yet been successfully captured and these neural changes may have prepared all four lobes for subsequent attentional capture.

### Frontal-posterior coupling for attentional capture: emergence and evolvement across time

Until the medium bins (**Fig.3G**), there appeared significant FP coupling (also for FF and FT pairs, attended condition; diamond symbols indicate absence of coupling in unattended condition) in response to visible versus invisible stimuli, and this frontal-posterior coupling persisted across medium bins (**Fig.3H**). In sharp contrast with the absence of coupling in the unattended state and early preconscious state, findings here highlight frontal-posterior coupling (especially FP coupling) as a neural marker of successful attentional capture. Based on findings of Exp.1 linking attentional capture and visual consciousness in our paradigm, we assume consciousness of the image must have been present during this intermediate stage.

No conscious experience can sustain indefinitely after stimulus offset. In the late bins as conscious experience finally faded, the posterior lobes (O, T, and P) first disentangled from each other (i.e., negative coherence), followed by FF and FT, and finally terminated by FP pairs (**Fig. 3G** and **3H**). These findings elucidate the dynamic nature of attentional capture process. Consistently, temporal generalization analysis also revealed time-evolving patterns of neural representations of consciousness along with time (**Fig.S11**). Altogether, findings here suggest that conscious experience of the image is accompanied by temporally varying large-scale functional connectivity patterns within hundreds of milliseconds.

Researchers asserting frontal-centric views of consciousness may treat frontal-posterior coupling as means of globally broadcasting real-time consciousness contents (*23*), feedback predictions (*25*) or higher-order meta-representations (*24*) across the brain, a process hypothesized to generate subjective experience. However, such possibilities of broadcasting fine-grained representations had been reexamined and opposed as a result (**Fig.4**).

**Figure 4.**
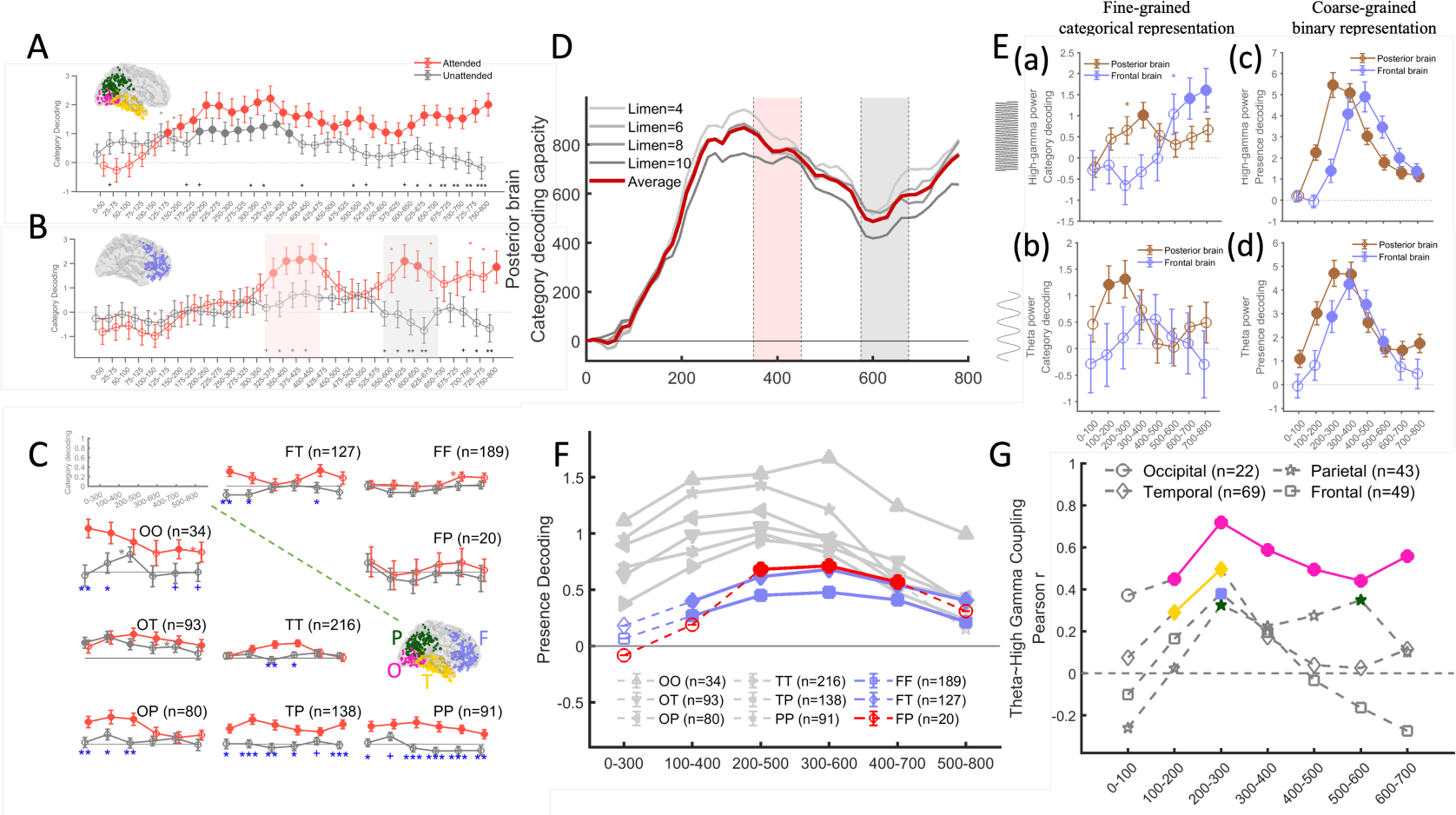
Persistently and globally exchanged signals are coarse-grained but not fine-grained representations of consciousness. (A-C) The posterior brain represented fine-grained categorical contents of consciousness in a persistent profile (A), as decoded by both regional activity and inter-lobe coherence measures (C, blue * denote significant difference between two conditions), while the frontal brain represented contents in a phasic profile (B). (D) During the initial period when the frontal brain started to represent contents, the summed amount of content representations of the posterior brain decreased rather than increased. This observation was robust irrespective of the inclusion criterion (i.e., Limen) determining how many percentages of responsive contacts exceeding the chance level (33.33%) were included into this analysis. (E) The frontal brain (light blue lines) decoded fine-grained categorical contents using high gamma oscillations (not until the late stage, panel a) rather than slow theta oscillations (panel b), although both oscillations durably represented coarse-grained binary signal of consciousness (panel c and d). (F) This binary presence signal was also represented by measures of inter-lobe theta coherence, globally and persistently. (G) Conveyed by theta oscillations, these globally exchanged binary signals might coordinate local regional activity via power-based theta and high-gamma cross-frequency coupling, especially in the occipital and parietal lobes. *, p < 0.05, uncorrected; filled circles, p < 0.05, FDR corrected.

### Distinct profiles of the inhibitory alpha-band activity in the posterior versus frontal brain

To aid conscious perception of specific image, we suppose attention may have helped in precisely determining the contents-to-be-perceived. Our finding that attention induces functional reorganization at the posterior brain agrees well with this deduction. Furthermore, previous research suggests attention also engages the inhibitory alpha band oscillation to access stored knowledge (*11*). Specifically, an early increase in alpha activity inhibits background noise first, allowing emerging signals to pop out, and then selects the popped-out signal for further amplification by decreasing alpha activity (*11*). Here, as measured by alpha band power, we found clear evidence of initial increase of alpha activity (0-300ms). Simultaneously, the early decrease in micro-saccade behavior (**Fig.S12**), which reflects attentional orienting (*40–42*), indicated that attention had been oriented to these emerging signals (*42, 43*). After that, the decreased alpha-band power (400-700ms, **Fig.3I**) provide direct neural evidence that these emerging signals had successfully captured attention, and the increased micro-saccade behavior and dilated pupil size (**Fig.S12** and **S13**; not confounded by horizontal or vertical eye gaze location, **Fig.S14**) provide further physiological evidence that these signals were truly amplified as a result at the late stage.

Notably, such a two-stage structure of alpha activity was only observed at the posterior brain and in the attended condition (**Fig. 3I, 3K** and **Fig.S15**). When we calculated the correlation between the two-stage modulation of alpha power and the mean capacity of content representation, we found significant correlations for the posterior brain (**Fig. 3J**) but nonsignificant either for the frontal brain or at the unattended condition (**Fig.S16**). These findings provide direct support for Klimesch’s viewpoint (*11*), implying that the two-stage alpha activity is engaged by attention to filter consciousness contents in the posterior brain. Simultaneously, this two-stage structure further highlights the 300ms as a cutting-edge timing point separating apart stages before and after attentional capture.

To complete the process of attentional capture, we propose the supply of attention cannot be interrupted. It is well known that the two frontal brain hemispheres compete for control of attention through mutual inhibition, each striving to direct attention towards its contralateral hemifield (*44, 45*). Therefore, disrupting their balance can bias the direction of attention. As found, in the frontal brain, inhibitory alpha power was consistently lateralized to the ipsilateral hemisphere (**Fig. 3K** and **Fig.S15**). This indicates that the ipsilateral frontal lobe was continuously suppressed, thus preventing attention from being distracted and stabilizing attention towards where the image was displayed. In the unattended condition when attentional focus was directed elsewhere, the peripheral images failed to induce this post-stimulus inhibitory alpha activity, thus resulting in a failure of attentional capture.

### Neural representations globally exchanged after frontal-posterior coupling is established

Almost all neuronal theories of consciousness agree that sensory analyses are primarily performed by the posterior brain. Indeed, the posterior brain encoded categorical contents using regional activity (**Fig. 4A**) and inter-regional connectivity (**Fig.4C**, lower left). Importantly, these content representations were continuously and durably maintained across time, aligning well with our smooth experience. Posterior-centric views of consciousness (*27, 30, 46*) typically attribute conscious experience to this sustained posterior activity.

In contrast, frontal-centric views of consciousness consider this posterior activity as necessary but not sufficient for consciousness (*23–25*). For example, the global neuronal workspace theory posits that consciousness arises from the frontal brain while broadcasting hybrid representations to widely distributed simple processors in the posterior brain (*23*). This hypothesis was rigorously tested here and challenged as a result.

We first examined the temporal profile of content representations in the frontal brain. Running content decoding analysis on regional activity measures, we found a phasic profile (**Fig.4B**) characterized by an intermediate silence period (450∼600ms) of which decoding performance decreased to non-significant (t(_56_)=1.04, p=0.303) relative to an early (350∼450ms, paired sample t-test, t(_56_)=-2.80, p=0.007) and a late (>600ms, t(_56_)=-2.24, p=0.029) surge of content representations in the frontal brain. This discrete profile was also observed via temporal generalization analysis which revealed two isolated clusters each shortly evolving across time (**Fig.S11**, right upper panel). Furthermore, decoding analysis on inter-regional connectivity measures at the theta band (*12*) (**Fig.4C**, upper right) revealed a similar phasic profile for frontal-temporal (FT, corrected p<0.05) and frontal-frontal pairs (FF, uncorrected p<0.05), while the frontal-parietal pair (FP), proposed to gate attentional supply, never encoded contents. In sharp contrast, the posterior brain exhibited a persistent profile (**Fig.4C**, lower left). These discrete content representations of the frontal brain do not align with the temporal continuity of our conscious experience. Instead, they better align with post-perceptual processes (*47, 48*) (e.g., decision and action), which may unfold either reflexively (possibly corresponding to the early surge) or be held under deliberate control (possibly corresponding to the late surge).

Another key proposal of the frontal view of consciousness suggests that the majority of posterior brain are blind to consciousness contents before receiving global broadcasts from the frontal brain (*3, 23–25*). If this is true, then, on the one hand, the total amount of content representations in the posterior brain should be significantly enriched immediately after the frontal brain starts broadcasting. However, as revealed, we instead found a general decrease after the early surge of frontal content representations (**Fig.4D**, 350∼450ms, same window outlined at **Fig.4B**). On the other hand, the frontal brain requires a messenger to broadcast contents to distant regions. Nonetheless, although frontal contents could be decoded (preferentially at a late stage) by high gamma oscillations (**Fig.4E**, panel a, light blue curve) associated with local spiking activity (*36, 37*), they could never be encoded by slow theta oscillations (**Fig.4E** panel b, note the contrast between the frontal versus posterior brain) subserving long-range communications (*8, 13, 49*), thus making them less likely to reach distant regions.

Recently, a growing GNWT hypothesis (*22*) has suggested that the frontal brain can still generate a conscious experience even when neural activity there is silenced (e.g., silent working memory (*50, 51*)). However, in reality, the frontal brain never halts its response midway in the attended condition (**Fig.3D** and **Fig.S17**). Instead, it actively and persistently maintains another type of coarse-grained representations, which signal the emergence of consciousness in a binary (1/0) manner, in parallel with the posterior brain (**Fig.3C** and **Fig.S17**) and also in both high-gamma and 4-Hz peaked theta oscillations (**Fig.4E**, panel c and d). In addition to regional activity, these binary signals are also represented by inter-lobe connectivity across the whole brain (**Fig.4F**), also in a persistent profile. This profile aligns well with the temporal continuity of our conscious experience.

These findings collectively highlight a frontal-posterior loop of visual consciousness. Critically, what are globally exchanged within this loop are coarse-grained signals of consciousness emergence rather than fine-grained consciousness contents. Serving possibly as global coordinators, these binary signals (insensitive to consciousness details) conveyed by theta oscillations may play functions in coordinating distant local activity (particularly in the occipital and parietal lobes) through power-based theta and high gamma cross-frequency coupling (**Fig.4G** and **Fig.S18**).

### General discussions

Bringing all elements together, we elucidate how goal-directed attention and captured attention play distinct but collaborative roles in visual consciousness in both the behavioral and neural level. When peripheral images were required to be neglected, we found visible relative to invisible images evoked sustained and temporally generalizing neural patterns for up to 450ms in the posterior brain, plus activation of a small area in the frontal brain. Then, when attention was directed towards these images, both regional activity and inter-regional connectivity were modulated before frontal-posterior coupling got established, even though this part of attention failed to predict image detectability. Later (>300ms), the frontal-posterior coupling got established and maintained but the signals exchanged did not inform fine-grained image contents but instead reflect success or failure of attentional capture. The decrease of the inhibitory alpha activity as well as the excitation of eye-related physiological response suggest attention had indeed been successfully captured by the emerging signal in the late stage. Psychophysical results show this part of captured attention proportionally predicted image detectability. These results attribute consciousness to attention-based coupling between the frontal and posterior brain as a whole, rather than activity of either part alone (for related debates, see (*22, 52–55*)).

First, consciousness cannot be solely attributed to the posterior brain. In the unattended condition, homogeneous neural activity generalizing across time for as long as 450ms duration was found in the posterior brain despite the absence of frontal-posterior coupling. If it really corresponds to a conscious experience, such sustained and homogeneous activity would be explicitly perceived rather than being too swift to be reportable(*30, 34*). Some others might alternatively argue that posterior activities are too weak to generate consciousness in the unattended condition and they might give rise to consciousness if strong enough for example when being attended. Following this logic, given goal-directed attention had enhanced posterior activities at an earlier 200-300ms bin to equivalent neural strength with that at the late 400-500ms bin, consciousness should have emerged at these early bins before attentional capture occurs. If true, the degree of goal-directed attention should proportionally correlate with image detectability, which however turned out not the case as we found in experiment 1. In fact, this correlation was only tied to captured attention instead. Collectively and alternatively, findings here favor the tripartite view that, before attentional capture occurs, posterior activity enters into a ‘preconscious’ buffer state beyond unconsciousness but still not conscious (*34*). This successive joint collaboration between goal-directed attention and captured attention thus finally results in one, not two, types of consciousness (*56, 57*) (opposing (*28*)).

Second, the frontal brain cannot yet support consciousness alone. Currently, a ‘global-workspace’ hypothesis is popular, proposing that consciousness works by broadcasting detailed consciousness contents from the ‘global workspace’ located at the frontal brain to distant rest regions to allow for cooperative processing by large collections of specialized systems such as memory, reasoning, planning, decision-making, and so on (*23, 58, 59*). By rigorously examining and opposing this hypothesis, findings here alternatively suggest that consciousness works by globally broadcasting coarse-grained signals of consciousness presence. During this process, the finding that goal-directed attention induced continuously unbalanced distribution of the inhibitory alpha-band activity between two frontal hemispheres suggests the frontal brain may continuously operate as a module of attention, detecting the emergence of salient signal since early on and maintaining attentional direction to that signal thereafter till to the late stage. Beyond that, its intermittent representation of highly detailed consciousness contents is better suited for post-perceptual processes (e.g., introspective reporting and preparation for actions) in line with task requirements (*47, 48, 60*). As a hypothesis for future investigation, we speculate that its relatively early and late content representations are configured to support reflexive and deliberate actions, respectively.

Altogether, findings here suggest the frontal brain and the posterior brain are both necessary for consciousness, but both insufficient by itself. Instead, consciousness is a product of these two, with their interaction forming a functional frontal-posterior loop to capture attention. The way this loop works, exchanging simple binary signals across the whole brain and accordingly coordinating distant local activities based on theta-high gamma cross frequency coupling, sheds insights on the neural dynamics of attentional capture in our paradigm. Potentially serving as a saliency map (*61*), these binary signals provide location to direct attention, thus capturing attention for the prominent signals in the posterior brain. Since the degree of captured attention proportionally correlates with visual detectability (**Fig.1C, 1G and S2**), we suggest that this loop structure navigating attention to contents might play a pivotal role by utilizing captured attention as a cost while translating corresponding contents into consciousness as a gain.

### A new model of consciousness

Based on these considerations and neural findings here, we term the early-stage effect of goal-directed attention as ‘attentional modulation’, whereas the late-stage effect which correlates with image detectability as ‘attentional integration’. With this division, we originally propose an attentional integration model of consciousness (AIM, **Fig.5**) which holds the view that consciousness requires attention and can only emerge till to the late stage (>300ms) after early signals successively capture attention (**Fig.5A**).

**Figure 5.**
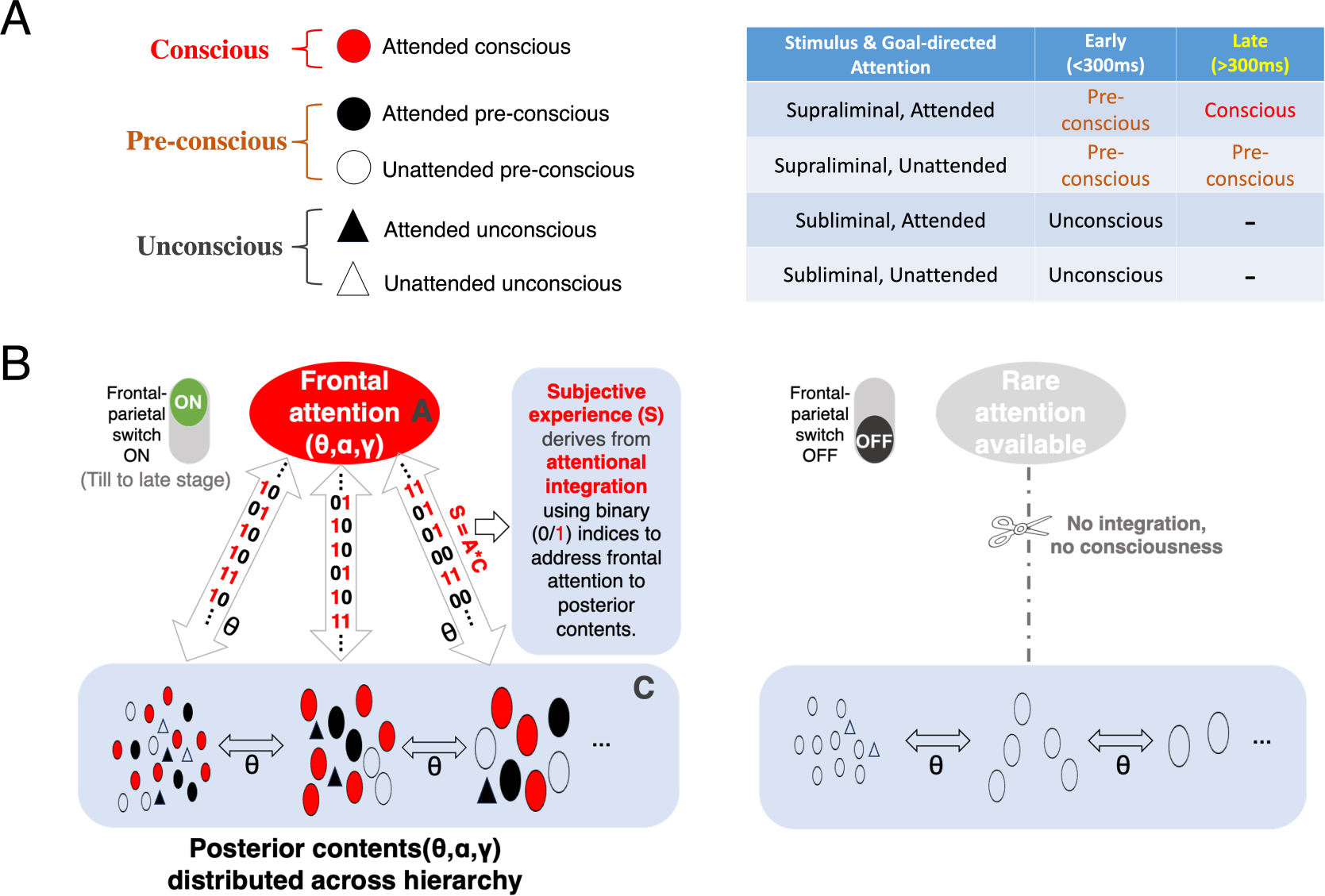
Attentional integration model (AIM) of consciousness. (A) AIM holds a tripartite view on the relationship between attention and consciousness. There is a preconscious buffer state inter-between unconscious and conscious (A, left panel). Consciousness can only emerge till to the late stage (>300ms) when preconscious signals get further integrated with captured attention (A, right panel). (B) in terms of neural mechanisms, AIM proposes that consciousness arises from an attention-content loop structure inter-between the frontal and posterior brain (B, left panel), where simple binary signals are exchanged to integrate frontal attention with posterior contents. In the context of visual consciousness in the human brain, contents are initially determined in the posterior brain, regardless of whether they are attended to or not. Subsequently, neural interactions (reflected by theta coherence measures) exchanging binary signals between the frontal and posterior brain enable these emerging contents (i.e., illustrated by red colored circles) distributed across cortical hierarchies to additionally capture more attention and thereby be consciously perceived. Conversely, failure in attentional integration (B, right panel) results in the absence of consciousness.

The fundamental idea of AIM can be encapsulated by a formula: S = A * C, suggesting that subjective consciousness (S) arises from the integration of attention (A) with contents (C) via an attention-content loop structure (**Fig.5B**). In terms of neural mechanisms of visual consciousness in the human brain (**Fig.5B**, left panel), this loop structure comprises the frontal brain, acting as a module for attentional control, and the posterior brain, serving as a module for representing the content stream. Inter-between them, attention-based frontal-posterior coupling is an essential prerequisite to consciously perceive external stimuli. Gated by FP coupling, their effective interaction would allocate frontal attention to posterior contents according to binary signals, thus completing the attentional integration process and transforming sensory contents into our conscious experience. If this interaction fails to occur (**Fig.5B**, right panel)—for instance, when the stimulus intensity is too faint (unconsciousness irrespective of attention), when attention is diverted elsewhere despite strong stimulus intensity (unattended preconsciousness), or when attentional capture process hasn’t commenced in time (attended early preconsciousness)—functional isolation between the frontal and posterior brain regions occurs. This isolation prevents frontal attention from accessing posterior contents, effectively reducing the amount of captured attention to zero and thus hindering the entry of visual consciousness.

AIM aligns partially with the Global Neuronal Workspace Theory (GNWT) (*23*) concerning the ‘ignition’ aspect, proposing that stimulus-induced frontal-parietal (FP) coupling serves as a gating mechanism for the interaction between the frontal and posterior brain regions. When an appropriate combination of goal-directed attention and stimulus-evoked posterior activity occurs, the FP switch is activated, establishing a loop between the frontal and posterior brain regions. However, where AIM notably diverges from GNWT is in how this frontal-posterior loop functions post-’ignition’. The primary distinction lies in the type of signals exchanged between the two regions. In AIM, the loop operates by exchanging coarse-grained signals directing attention rather than fine-grained signals representing consciousness contents. This observation suggests that a frontal ‘workspace’ (although possibly required for post-perceptual processes like response to stimulus) is not indispensable for consciousness per se. Though AIM is open to the idea that attention may ‘bind’ sensory features into a bound representation of objects (*62, 63*), or route multiple source information to the frontal brain and consolidated there into a convergent representation and then go back to other regions (*23, 59*), AIM does not confine consciousness to such convergent representations. Instead, AIM holds that the distributed representations at various levels of the sensory hierarchy can be consciously perceived if only integrated with captured attention. Put simply, AIM proposes consciousness arises from the integration process coupling each distributed representation with attention though not necessarily requiring low-level distributed representations to be bound into a higher-level hybrid one.

In the realm of consciousness research, various neuronal models have emerged over the past decades. Alongside the notion of global broadcasting (*23*), these models encompass ideas such as local recurrent processing (*30*), higher-order representation (*24*), top-down prediction (*25*), causal-effect loop structure (*27*), and attentional control (*64*). Many of these concepts can be discerned within the attention-content loop structure of AIM, albeit from distinct perspectives, yet none present a fully comprehensive view. For instance, the attention-content loop can also be construed as a causal-effect loop structure proposed by the integrated information theory (*27*), where each module can causally influence itself through interaction with the other. The distinction lies in the spatial scales at which these loops for consciousness operate-either encompassing both the frontal and posterior parts or restricted within the posterior brain alone. By amalgamating these disparate concepts, AIM stands poised to reconcile these sharp discrepancies in the field.

This study is not without its limitations. One constraint is our inability to precisely pinpoint the moment at which consciousness emerges in the attended condition. While we interpret frontal-parietal (FP) coupling as a contemporary indicator dissociating early preconscious from late conscious states, this aspect warrants more rigorous investigation in the future. Another limitation pertains to the absence of causality. Experimental interventions targeting the three components of AIM—accessible contents, goal-directed attention, and attentional capture—are necessary to empirically test this model in a causal manner. Lastly, it remains uncertain whether the current findings can be generalized to other paradigms and other conscious experiences, for example, involving more abstract content such as goals and volition. In such cases, given that the frontal brain would act as a component of both attention and content modules (*47, 48, 65*), future studies are awaited to elucidate whether the AIM loop structure can persist within the frontal brain or if it necessitates alternative, yet undiscovered mechanisms. In the current era of exploring conscious artificial intelligence agents (*66, 67*), the attention-content loop proposed for attentional integration may also inspire novel engineering approaches.

## Supporting information

Movie S1. Demo illustration of trials without any probes displayed in the pretest session (number-counting task) of Exp.1.

Movie S2. Demo illustration of trials with randomly selected occasional probes displayed in the posttest session (number-counting task) of Exp.1.

Movie S3. Demo illustration of trials displayed in the unattended blocks (number-counting task) of Exp.2.

Movie S4. Demo illustration of trials displayed in the attended blocks (image-categorization task) of Exp.2.

Movie S5. Spatial distribution of responsive contacts across four neocortex lobes and across time.

## Acknowledgements

We give thanks to Hakwan Lau, Chencan Qian, Qing Yu and Xiangyong Yuan for insightful comments on earlier version manuscript, to Zhihao Guo and Bingqing Zhang for help in iEEG data collection, and to Haiyan Kong and Chenyu Gao for help in behavior data collection.

## Author contributions

Conceptualization: XZ, LW, YJ

Methodology: XZ, LW, YJ, KZ

Investigation: XZ, CZ, XW, WZ

Visualization: XZ, LW, CZ, KZ

Funding acquisition: LW, SH, YJ, XZ

Project administration: LW, KZ, XZ

Resources: YJ, KZ, LW

Supervision: LW, KZ

Theoretical modelling: XZ

Writing – original draft: XZ

Writing – review & editing: XZ, LW, SH

## Materials and Methods

### Participants

Twenty-one healthy participants (16 females, 24.19±2.54 years) took part in Experiment 1. For two participants, audio data recording oral reports for probe trials in the posttest session were not correctly recorded due to technical issue. These two participants were thus excluded from analysis of oral reports but were included in the analyses of counting performance. To replicate results of Exp 1, a follow-up experiment (more details see Fig.S2) was designed and performed using roughly the same paradigm and comparable sample size (20 healthy colleague students, 16 females, age=23.1 ± 1.88 years) but recruiting a different experimenter and two new audio encoders.

Twenty-one drug-resistant epilepsy patients (10 females, 24.48±6.91 years) participated in Experiment 2. Nineteen patients were from Tiantan Hospital, Capital Medical University, and two patients were from Yuquan Hospital, Tsinghua University. These patients underwent intracranial electrode implantation as part of their clinical evaluation for epilepsy surgery when noninvasive studies could not adequately localize the origin of their seizures. Among them, six patients did not have their pupil data recorded: the first four (S01-04) due to the eye tracker not being in place, and the other two (S16 and S17) due to poor eye signal quality, which prevented successful eye tracking. These six patients were excluded from pupil data analysis but were included in other analyses.

For both Experiment 1 and 2, all participants had normal or corrected-to-normal vision, had no history of head trauma or encephalitis, and provided written informed consent. The research protocol of Experiment 1 was approved by the review board at the Institute of Psychology, Chinese Academy of Sciences, and that of Experiment 2 was approved by the institutional review boards at the hospital sites.

### Apparatus

The stimuli were presented using Psychophysics Toolbox (*68, 69*) for MATLAB (The Mathworks, Natick, MA, USA) on a 21-inch gamma-corrected monitor with a resolution of 1920 × 1020 pixels and a refresh rate of 60 Hz. An infrared eye tracker system Eyelink Portable Duo (SR research, Kanata, Ontario, Canada) was used to monitor gaze location, (micro)saccade behavior and pupil size in both Exp1 and 2. Observers viewed the screen from about 60 cm away, using a chin rest to stabilize the head position in Exp1. The chin rest was not used in Exp 2, where, instead, epilepsy patients sitting in bed were told to minimize their head movement during the recording period (in remote recording mode).

### Materials

Visual stimuli used consisted of central and peripheral streams. The central visual stream consisted of four numerical values (4, 5, 6, or 7) which were employed to create a rapidly alternating number sequence at a frequency of 10 Hz, presented within the central visual field.

The peripheral visual stream consisted of 48 visual stimuli, comprising 16 animal images, 16 object images, and 16 face images, and were presented at 4.0° visual eccentricity (Fig.1A). Additionally, eight visual masks originally constructed using line segments and then created into different variants by rotating and flipping this original mask were included in the peripheral visual stream. A pair of nonmatching masks was randomly selected for each trial. The animal and object images, as well as the visual masks, were sourced from (*70*). The face images which are all emotionally neutral and differ in gender half by half, were initially obtained from (*71, 72*) and were then converted into simple line drawings using an open-source tool (http://jianbihua.renrensousuo.com/).

Both central and peripheral streams were controlled using Psychophysics Toolbox version 3 for MATLAB. All stimuli were presented on a white background (1920 × 1080 pixels, 60Hz monitor; thinkpad T490). Participants were seated 60cm from the monitor, thus making the average size of the line drawings 4.8° in side length, and the masks 6°.

### Paradigm design and procedure

In the experiment, participants were instructed to fixate on a central cross while a central fast-alternating number stream and a peripheral visual stream were presented simultaneously and unfolding across time in each own pace. Participants maintained fixation on a small black fixation dot (0.5° in diameter) that was continuously visible throughout the procedure. Four demo videos of the stimulus presentation sequences are available in the Supplementary materials.

### Central number sequence

For each trial, the central visual stream was a pseudo-randomly ordered temporal sequence of single number characters altering at 10 Hz. For number-counting blocks, the target number to be counted was chosen randomly and informed to the participant at the beginning of each block.

### Peripheral visual stream consisting of masks and image

For each block of 72 trials, forty-eight images were divided into four groups (A, B, C, and D), each containing four animals, four objects, and four faces, with images from only one group used in any given block. Within a block, 24 visible image trials (twice per image), 24 invisible image trials (twice per image), 12 visible control trials and 12 invisible control trials were conducted in random order. All trials were conducted in mixed and random order. There was a 1000ms blank intertrial interval that occurred after each trial and before the next.

Visibility manipulation closely followed the methodology outlined in (*70, 73*). Visible trials in the peripheral stream comprised a ‘mask-blank-image-blank-mask’ sequence, with each mask and blank period lasting 200ms and 100ms, respectively, resulting in a total trial duration of 633ms. The image was surrounded by blanks, making it weakly masked and explicitly visible. In contrast, invisible trials followed a ‘blank-mask-image-mask-blank’ sequence, with the image closely surrounded by masks, rendering it heavily masked and thus invisible to observers. Both visible and invisible trials were followed by a 1000ms blank intertrial interval before the next trial.

For both visible and invisible trials, their control trials were set to mimic the mask and blank sequence, but replaced the middle image with a complete blank of the same duration (33ms). We subtracted different measures of interest (see following) of control trials from that of invisible and visible trials (exampled at Fig.2A), obtaining a clean mask-free measure tied to visibility state when the image was either invisible or visible, respectively.

### General procedure of Exp1 and Exp2

Both Experiment 1 and Experiment 2 were conducted using a block design. In all trials and blocks, the central number stream was consistently presented alongside the peripheral image and mask stream. We manipulated attentional focus across different conditions: directing attention primarily to the central number stream (pretest of Exp1 and unattended blocks of Exp2), focusing mostly on the central number stream but also partially on the peripheral stream (posttest of Exp1, achieved by randomly adding occasional probes), or directing attention primarily to the peripheral stream (attended block of Exp2).

In the pretest session of Experiment 1 (i.e., without probes, consisting of 2 blocks), participants were instructed to focus solely on counting while disregarding the peripheral stream. To achieve so, participants were not informed about the posttest session, which included probes, until after the pretest session. The peripheral stream was presented in either the left or right visual field, with the order determined randomly for each trial, but no more than three successive trials occurred at the same side.

In the posttest session of Exp1 (i.e., with probe, 10 blocks), we still required participants to set high priority the counting task and perform at their best. The only difference was that this session shifted a bit of attention to peripheral image, by occasionally probing its visibility in 16% randomly selected trials (**Fig.1D**, Method). In these trials, a probe cue, i.e., a square outlining the image (side length 4.8°), interrupted the trial sequence at two timepoints (0ms or 200ms after image offset). We set 12 probes per block, with any two probes interleaved by 4∼8 trials though participants were not informed of this rule. These probes appeared in 4 visible image trials, 4 invisible image trials, 2 visible control trials and another 2 invisible control trials. Half probes appeared at 0ms after image offset, half at 200ms after image offset.

When seeing this probe, participants first pressed keys to complete counting as usual, to indicate how many target numbers they saw before probe. This was designed to hold the majority of attention towards counting, performing the counting task as best as possible while leaving only a small portion distracted to peripheral images. Otherwise, if too much attention was distracted to image, wo proposed too much attention supplied to image would lead to a saturation ceiling effect hindering us to detect to what extent consciousness might decay within a 200ms temporal gap (200ms versus 0ms). After the first key-press, color of the probe cue turned red, indicating the start of audio recording. Seeing this, participants were required to orally reported whether they saw the target image surrounded by masks. If yes, further reported its category and detailed identity. A typical report for probe within a visible image trial was like ‘I am sure I saw an animal image with a tail but I am not sure whether it was a cat or a dog’.

In experiment 2 (block design), attention was directed to different streams randomly alternating block by block but with no more than two repetitions, to peripheral image in attended blocks but to central counting stream in unattended blocks. In attended blocks of Exp2, participants focused on an image categorization task, but neglecting the central number stream completely. To help stabilize attention to peripheral image, peripheral stream was displayed consistently in one visual field (either left or right) for all trials but randomly set across blocks. At the end of each trial, participant was provided a forced choice to class the peripheral critical image into either animal, object or face by pressing a corresponding key. In attended blocks, counting task was not performed any more. In unattended blocks of Exp2, same stimulus streams were presented on the screen and the only difference was that participants performed the central counting task instead. The image categorization task was not performed anymore and participants were required to totally ignore the peripheral stream.

### Audio data collection and offline coding

Audio data were collected for probe trials during the posttest session of Experiment 1. The laptop’s built-in recording device was activated immediately after participants pressed the response key which indicated when they saw the target number. Then, participants orally reported whether they perceived the critical image (excluding masks) at the onset of the probe cue, if so, they identified the category and identity of the image. Audio data were recorded in real-time and later analyzed offline by two encoders. Importantly, both coders were blinded to the experimental conditions to ensure unbiased coding.

The audio data were coded into three dimensions. In the first dimension of visibility (i.e., whether the critical image was seen), scores of 0-3 corresponded to surely not seen, seemingly not seen, seemingly seen, and surely seen, respectively. In the dimensions of categorical and item-level accuracy (i.e., whether the reported category/item was correct), scores of 0-4 corresponded to surely wrong, likely wrong, neutrally not sure, likely correct, and surely correct, respectively. Between two coders, interrater reliability was high in all three dimensions (correlation coefficient close to 1, Fig.S1A).

### Pupil data collection

During Experiment 1, participants rested their heads on a chin-rest to limit head movement. In Experiment 2, the chin-rest constraint was removed to increase comfort for epilepsy patients who were sitting in bed. However, participants in both experiments were instructed to minimize head movements during the recording period. In both experiments, eye-tracking data were acquired using the EyeLink Portable Duo SR Research eye-tracker with 1 ms temporal resolution (at 1000 Hz frequency) in remote head-free-to-move mode, recording data from the monocular eye with a higher signal-to-noise ratio. The camera-to-eye distance roughly fluctuated around 60 cm.

### Pupil data preprocessing

The raw sampling data was preprocessed using the Matlab interface to Eyelink EDF files (https://osf.io/fxumn/, Version 1.0.1) provided by SR research to identify eye-tracking indices. Relevant eye-tracking events, such as fixations, blinks, and saccades (especially microsaccades), were detected for each participant. This was achieved using the functions edfExtractInterestingEvents.m and edfExtractMicrosaccades.m. The relevant variable values and timing information were then extracted using edfExtractVariables.m and edfExtractKeyEventsTiming.m, respectively.

For each participant, time series of eye-related variables (pupil size and microsaccade ratio, see following) were calculated independently for each combination of attention (attended versus unattended) and visibility (visible versus invisible, as well as corresponding control trials). This was based on epochs lasting from 500 ms before image onset to 1500 ms after image onset. Trials with missing values (typically caused by eye blinks) exceeding 25% of the timepoints were discarded from subsequent analysis.

### Pupil data analysis

**Pupil Size.** Following the protocol provided by SR Research (version 1.0), we utilized an artificial eye to convert arbitrary pupil size units to millimeters. This process involved calculating a scaling factor (0.106) based on a distance parameter set at 60 cm. To convert pupil data from arbitrary units to diameter, we multiplied the square root of the pupil area data by the scaling factor (0.106). For instance, for a typical pupil area data value of 1216, which represents the average mean of trials from our first recording block, the corresponding pupil diameter would be sqrt (1216) * 0.106 = 3.70 mm.

It’s important to note that this scaling factor is tied to the distance parameter and would change if the distance parameter were altered. Additionally, pupil data in both experiments was recorded in a head-free-to-move mode (i.e., remote mode), which could lead to fluctuations in the distance parameter. Although we reminded participants to maintain the ideal distance setting (60 cm), especially in Experiment 2 where a chin rest was not used, these fluctuations may have been more pronounced. To this end, all subsequent analyses on pupil size were conducted using the pupil area data in arbitrary units.

### Microsaccade detection and analysis

Microsaccade events were extracted from the raw eye position data using the algorithm described in Engbert & Kliegl (2003), implemented with default settings in edfExtractMicrosaccades.m. Velocity was computed using a time span of 20 ms around the current sample. The onset of a microsaccade event was defined as the time point at which velocity exceeded the trial’s median velocity by 6 or more standard deviations. The minimal microsaccade duration was set to 12 ms, and the minimal time allowed between two microsaccades was also set to 12 ms; otherwise, they were merged into a single event spanning from the onset of the first microsaccade to the offset of the second one. This approach enabled the detection of microsaccade events for every trial. Timepoints corresponding to microsaccade events were coded as 1, while other timepoints were coded as 0.

The microsaccade ratio was then calculated as the percentage of trials containing a microsaccade event for each timepoint (1 ms), resulting in a percentage time course. This time course was smoothed using a 100 ms moving window (following (*42*)), replacing each center value by averaging values within its window. The smoothed data was then detrended to remove baseline drift or slow trends from signals using a polynomial detrending algorithm with the break point set to image onset.

### iEEG: electrode/contact reconstruction and localization

Postoperative computed tomography (CT) images were coregistered to preoperative T1-weighted MR images using FreeSurfer (v6.0.0, http://surfer.nmr.mgh.harvard.edu/). The registration was visually verified and manually adjusted if necessary. The implanted electrodes and contacts were reconstructed using the stereotactic localization software (*74*).

Initially, the intracranial electroencephalography (iEEG) contacts were labeled with a cortical area based on Freesurfer’s anatomical parcellation (*75*) in native space. Subsequently, all converted iEEG contacts were further labeled in standard space using the Desikan-Killiany (DK) labeling protocol (*76*), which is a widely used human cortical labeling protocol. During this process, the coordinates of all contacts were mapped onto a standard MNI space. Based on the MNI coordinates, contacts were further labeled to functional parcellations using atlases like the Human Connectome Atlas (*77*) and its extension (*78*), as well as the Yeo 7 network (*79*), to evaluate their functional roles when necessary.

To distinguish gray matter (GM) contacts from white matter (WM) ones, a spherical region of interest (ROI) with a radius of 3.5 mm was first created for each contact around the electrode coordinate in native space and then transformed to the standard space. The percentage of voxels labeled as GM within the ROIs (inclusion) was calculated and compared to the percentage of voxels outside the ROIs (exclusion). If the probability of inclusion exceeded that of exclusion, the ROI was classified as GM; otherwise, it was categorized as WM or an unknown area. For each included ROI, the probability values were compared between brain areas involved, and the ROI contact was labeled as the structural area with the maximal probability value.

### iEEG: data collection and preprocessing

For all 21 patients, intracranial electroencephalography (iEEG) data were recorded using a Nihon-Kohden recording system at a sampling rate of 2000 Hz. To ensure data quality and avoid potential confounds from interictal activity or seizures, we verified that no subclinical or clinical seizures occurred during or immediately before the experiment.

After acquisition, the iEEG data were analyzed using MATLAB combined with EEGLAB toolbox. The data were first bandpass-filtered from 0.5 to 200 Hz using a zero-phase delay finite impulse response filter and then downsampled to 500 Hz. Noise at 50 Hz and its harmonics were removed using a notch filter. Subsequently, iEEG traces from each contact were visually inspected to identify and exclude bad channels and time segments containing artifacts such as epileptic activity, large-amplitude slow-wave drifts, or high-frequency activity. Importantly, this visual inspection was performed in a blinded manner to the experimental conditions. For each patient, the time series of all remaining valid contacts were re-referenced by subtracting the mean signal of all white matter (WM) contacts. This step aimed to remove globally shared background signals. This unbiased approach avoids introducing inter-contact variance due to the re-referencing procedure and enhances the replicability of the findings.

### iEEG: ERP analysis

For event-related potential (ERP) analysis, preprocessed traces were first bandpass-filtered between 0.5 and 30 Hz before segmenting epochs. Since band filtering usually introduces edge artifacts at the head and tail of traces, this filtering step was crucial to ensure that data inter-between the head and tail (including all data of interest) were not contaminated by these artifacts. After filtering, the traces were segmented into epochs spanning from −500 ms before to 1000 ms after image onset. Subsequently, outliers in the remaining data (time * epochs) of each participant were identified and replaced by NaNs if their voltage value surpassed 5 standard deviations of all epochs at any time point. After baseline correction (200ms pre-stimulus baseline), the epoch data were submitted to statistical analyses. Note that these processes were performed upon non-NaN elements, while treating NaNs as missing values.

### iEEG: time-frequency decomposition

For time-frequency analysis (TFA), we first bandpass-filtered the preprocessed traces between 0.5 and 200 Hz before segmenting epochs. Prior to epoch segmentation, outliers within each trace were identified and replaced with NaNs, following the same procedure as the event-related potential (ERP) analysis. After that, NaN values within traces were interpolated using the inpaint_nans.m function (https://www.mathworks.com/matlabcentral/fileexchange/4551-inpaint_nans), which uses a simple plate metaphor, before time-frequency decomposition.

The TFA decomposition was conducted using Morlet wavelets via the newtimef.m function of the EEGLAB toolbox. The length of the wavelet increased linearly from 3 cycles at 2 Hz to 54 cycles at 180 Hz, corresponding to a sliding window length of 1670 ms. This modified wavelet length was chosen to optimize the tradeoff between temporal resolution at lower frequencies and stability at higher frequencies. The TFA outputted complex values combining both amplitude and phase values for each time-frequency point, resulting in a 2-D plane with a log-spaced frequency dimension (60 bins from 2 to 180 Hz) and a linearly spaced time dimension sampled at 250 Hz (4ms resolution). In the dimension of time, this 2-D plane was segmented into epochs ranging from −600ms to 1000ms around image onset.

### iEEG: neural oscillation power

For each complex value at every time-frequency point, we extracted the amplitude and calculated the power as the squared amplitude. The power values of each epoch were then z-normalized across time to reduce sensitivity to noisy trials and allow for comparison of power values across different frequency bins. Subsequently, every normalized power epoch was baseline corrected using a 200-ms pre-stimulus baseline. Finally, the power values were averaged across frequencies to assess the change in mean band power following image onset for any frequency bands of interest. It is important to note that mask-evoked power change was specifically measured by control trials, which could have been removed via the contrast between image (masks plus image) versus control (masks only) trials, resulting in a clear measure tied to visible or invisible processing in either the attended or unattended condition.

To test whether inhibitory alpha band power during the pre-stimulus period was selectively allocated to the hemisphere ipsilateral to the peripheral visual stream, we contrasted contralateral versus ipsilateral trials on its oscillation power for each contact located in either the left or right hemisphere. Baseline removal and normalization across time were not conducted anymore. Instead, power values were normalized across epochs for each time point to scale the data while preserving the variance between trials. Since visual masks and the target image were arranged in different timing sequences between visible versus invisible trials (but in the same sequence within each), which might differently alter the temporal profile of alpha power activity, we performed this analysis in parallel for visible trials and invisible trials, respectively.

### iEEG: mutual distance between contacts

To assess the spatial clustering of responsive contacts in each lobe, particularly for the unattended condition, we computed the mutual distance between any two responsive contacts based on their MNI coordinates. To reduce variability, we first flipped the MNI coordinates of right hemisphere contacts to symmetric locations at the left hemisphere. Then, Euclidean distance was calculated for any two contacts. The mutual distance was finally calculated as the mean of all pairwise Euclidean distance.

Next, we performed a surrogate procedure to create a chance distribution of mutual distances. For each electrode, including the one with the maximum number of responsive contacts, we randomly replaced it with another electrode that contained an equivalent or greater number of contacts within the same lobe. From these contacts, an equal number of contacts were randomly selected, even if this included non-responsive contacts. Each electrode and contact could only be selected once at most for each surrogate. By repeating this procedure 10,000 times, we generated a chance-level distribution of mutual distances. This distribution was used to rank the true distance value, providing an indication of the spatial clustering of responsive contacts relative to chance.

### iEEG: functional connectivity between contact pairs

Between any two responsive contacts, functional connectivity was calculated across time rather than across trials. This allowed us to assess the degree of phase clustering for each condition without necessarily requiring phase differences to cluster around the same angles across trials. More specifically, an approach based on imaginary coherence (*80*), which is insensitive to volume conduction, was employed to mitigate the impact of a common input on interregional neural synchronization (*81, 82*). Using code from (*83*), imaginary coherence for each frequency was calculated as:

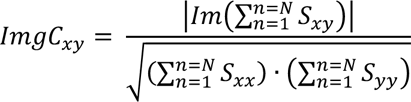

where S_xy_ denotes the cross-spectral density between activities at contacts x and y, S_xx_ and S_yy_ are the autospectral densities for contacts x and y, N denotes the number of time bins, and ImgC represents the absolute imaginary coherence. Cross- and autospectral densities were computed using time series of complex values obtained from Morlet wavelets introduced above (iEEG: time-frequency decomposition).

For each trial, ImgC was computed between any two contacts over 300 ms time segments in a sliding manner. The resulting values were then averaged across corresponding trials to obtain coherence measures for each combined condition across attention and visibility. In this study, six sliding time segments of a 300 ms window size were used: 0-300 ms, 100-400 ms, 200-500 ms, 300-600 ms, 400-700 ms, and 500-800 ms, to illustrate how coherence measures evolved over time.

The selection of time segment length involves a trade-off between signal-to-noise ratio and temporal precision. Longer time segments, encompassing more oscillation cycles, yield more robust and higher signal-to-noise estimates of phase synchronization. However, this comes at the expense of decreased temporal precision for time-varying and task-related modulations of phase synchronization, which are the main focus of this study. To reduce susceptibility to noise or nonrepresentative data, the resulting ImgC value for each condition was transformed to adjusted ImgC (Rayleigh’s Z, using code from (*83*)) using the formula:

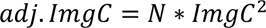

where N is the number of timepoints. This calculation allows the resulting coherence values to exceed 1. A p-value can be computed from adj.ImgC to determine statistical significance (*83*).

### iEEG: decoding analysis

The decoding methodology typically involved three main components: a classifier, label data and feature data, with the aim of classifying feature data into different labels. In terms of classifiers, classifiers used in the present study were Support Vector Machine (SVM) classifiers with a linear kernel. Without otherwise delineated, classifiers were typically trained in a sliding bin manner (bin size = 25ms, step size = 15ms), with 10ms overlap between two neighboring bins. In this way, models were trained independently for each contact and each bin.

Without otherwise delineated, the feature data typically used for decoding were multi-variate, consisting of mean ERP response and the mean normalized power of six contiguous frequency bands (2-4, 4-8, 8-13, 13-30, 30-60, and 60-150 Hz), with all averaged within the bin. In other scenarios, feature data could also consist of some univariate measures (e.g., theta power or high gamma power), including phase coherence measures calculated for each trial, to better evaluate contribution of specific features.

Setting of the label data reflect our design of interest. Basically, two types of labels were trained: one type of coarse-grained representation of image presence and another one of fine-grained representation of categorical content. The former (coarse-grained) utilized all trials, including both image trials (labeled as ‘image present’) and control trials (labeled as ‘image absent’). Models were trained to predict whether an image had presented (chance level=1/2) under each combined condition of attention and visibility. Since the number of image trials was double that of control trials, each training was split into two runs, with each run using non-overlapping half of the image trials to distinguish from the control trials, and their decoding performance was averaged across two runs. In each run, an 8-fold cross-validation approach was utilized, where the feature data were divided into eight equivalent datasets. In each fold, seven datasets were used for training the classifier, while the remaining dataset was used for testing. This process was repeated for all eight folds, allowing all samples to be tested once. The performance of the classifiers was evaluated based on decoding accuracy, calculated as the ratio of correctly classified samples to the total number of samples.

By contrast, fine-grained decoding of categorical content was performed by utilizing only image trials (control trials were not used since no images were presented) with the aim to decode pattern difference between three image categories (chance level=1/3). Since different labels were of equal trial number, model training was performed in one run by utilizing the same 8-fold cross-validation approach. Finally, for each of four combined conditions of attention and visibility, decoding accuracy for the fine-grained representations was also calculated as the ratio of correctly classified samples to the total number of samples.

### Visibility-related measures

Activities associated with either invisible or visible processing of images were calculated by subtracting control trials (masks only) from corresponding image trials (masks and image), obtaining a mask-free measure linked to each condition. Then, a visibility index was quantified using the following formula.

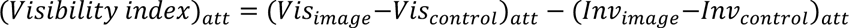

Here, “Vis” and “Inv” denote the mean value of measures of interest in the visible and invisible conditions, respectively, while the resulting visibility index quantifies their difference and “att” denotes the attended or unattended condition for each. Though this visibility-related measure had been used to evaluate neural correlates of consciousness (NCC) (*70, 73*), we were aware that this index cannot be interpreted as a clear measure of NCC due to its confounding with factors like stimulus intensity and performance capacity(*84*). Our main interest was to calculate the contrast between two conditions (attended minus unattended), so as to evaluate how goal-directed attention alters visibility-related neural activity for each responsive contact and time bin. Without otherwise specified, all dependent variables (i.e., various measures about power, coherence, and decoding performance) reported in the manuscript and depicted within figures had been calculated using this visibility index algorithm.

Taking decoding performance as an example. To decode for both coarse-grained and fine-grained representation of consciousness-related signal, respectively, we first trained SVM models for each combined condition of attention and visibility: attended visible, attended invisible, unattended visible and unattended invisible. Importing the decoding performance to the visibility index algorithm, we then obtained visibility-related decoding performance (visible-invisible) for each type of neural representations for each contact and time bin. This allowed us to evaluate how visibility representations evolved in the dimension of space and time. Replicating this procedure for attended and unattended condition, independently, and further making a contrast between them (attended versus unattended), we finally got to evaluate the impacts of attention on both types of visibility-related neural representations.

### Responsive contacts

In the current era, a hotly debated topic regarding the mechanisms of consciousness (*22, 54*) revolves around the functional differences between the anterior brain, particularly the frontal lobe, and the posterior brain, encompassing the occipital, temporal, and parietal lobes. Until this issue is resolved, elucidating the roles of more specific brain regions remains challenging. Hence, in this study, responsive contacts were defined as contacts that are responsive to consciousness-related signal in either attended or unattended condition. More specifically, they were selected from the gray matter of the four neocortical lobes (Occipital, Temporal, Parietal and Frontal lobes), while excluding those from white matter. In terms of neural activity, responsive contacts were identified based on either event-related potential (ERP) responses contributed mainly by slow neural oscillations (here filtered between 0.5-30 Hz) or fast gamma oscillation (70-150 Hz) power responses whose activity correlates with local neuronal spiking (*36, 37*).

Taking the ERP activity as an example first. For each attention condition and timepoint, a true NCC measure were calculated using the Visibility algorithm introduced above. Subsequently, the visibility labels of epochs were randomly shuffled while keeping block membership and attention label unchanged, and NCC measures were recomputed using the same formula. This process was repeated 10,000 times to create a null distribution for each timepoint. The true NCC measure was then ranked within the 10,000 surrogate distribution, with ranks of >=9995 or <=5 considered significant (corresponding to a two-tailed P <= 0.001). Timepoints showing significant activity were clustered across time, with clusters lasting >33ms considered responsive. Timepoints inside the identified clusters were coded as 1 while other timepoints outside as 0.

The same procedure was conducted for high gamma power in parallel with ERP analysis, resulting in another timeline of binary indices. For tach timepoint, above-chance contacts were defined as the union of contacts that turned out above-chance for either ERP or high gamma power analysis.

In the second level analysis, for each timepoint, we summed the number of above-chance contacts from the four neocortex lobes of interest, resulting in a new timeline of summed N. To control for false alarms, we calculated the mean value (*n*) and standard deviation (*std*) across timepoints during the 500ms pre-stimulus baseline period. We then calculated a Z value for each timepoint using the formula:

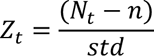

where N_t_ denotes the number of above-chance contacts for each timepoint. Responsive window included timepoints of a Z value >3 and falling within the range of [0 600]ms post-stimulus onset. Responsive window for the attended condition turned out much exceeding that for the unattended condition. For each attended and unattended condition, respectively, responsive contacts were identified as the union of above-chance contacts derived from all timepoints within the responsive window.

### Responsive units reducing data homogeneity

For each lobe, patients may have multiple electrodes implanted, with each electrode potentially containing multiple responsive contacts. It is well known that data of adjacent contacts coming from the same electrode are highly correlated. On the contrary, data of contacts from the same electrode but different lobes, as well as those from different electrodes but the same lobe, are much more heterogeneous. We thus conducted second-level statistical analyses by considering each responsive unit as an independent sample. Specifically, data from n responsive contacts coming from the same lobe and same electrode were averaged to represent the unit activity. To further mitigate contamination from false alarm noise, unless specified otherwise, we only included units that contained n>=2 responsive contacts into the second-level analysis of regional power and decoding performance.

Similarly, imaginary coherence values were also computed at the unit level, with unit-level coherence calculated as the mean of all included contact-level pairs. To control for false alarm, unless specified otherwise, we only included unit pairs that contained n>=4 contact pairs into the second-level analysis of inter-unit coherence.

### Statistics

Considering each responsive unit as an independent sample for second-level analyses, parametric tests used included one-way and two-way repeated measures analysis of variance (ANOVA), paired-sample t-test, one-sample t-test, as well as Pearson and Spearman correlation analysis. All statistical tests were conducted in a two-tailed manner with a threshold of p = 0.05. The FDR correction or cluster-based permutation test was used to correct for multiple comparisons with a threshold of corrected p = 0.05.

To replicate findings, we also conducted linear mixed effects (LME) models with conditions of interest (e.g., visibility and/or attention) as fixed effects, and with patient and electrode membership (not individual contacts) as random effects. Using the restricted maximum likelihood method to estimate LME parameters, and using Bonferroni correction to correct for multiple comparisons (corrected P=0.05), the LME-based approach, which requires significantly more calculations, resulted in similar findings (not shown here for simplicity) as those reported here.

---

**Fig.S1.**
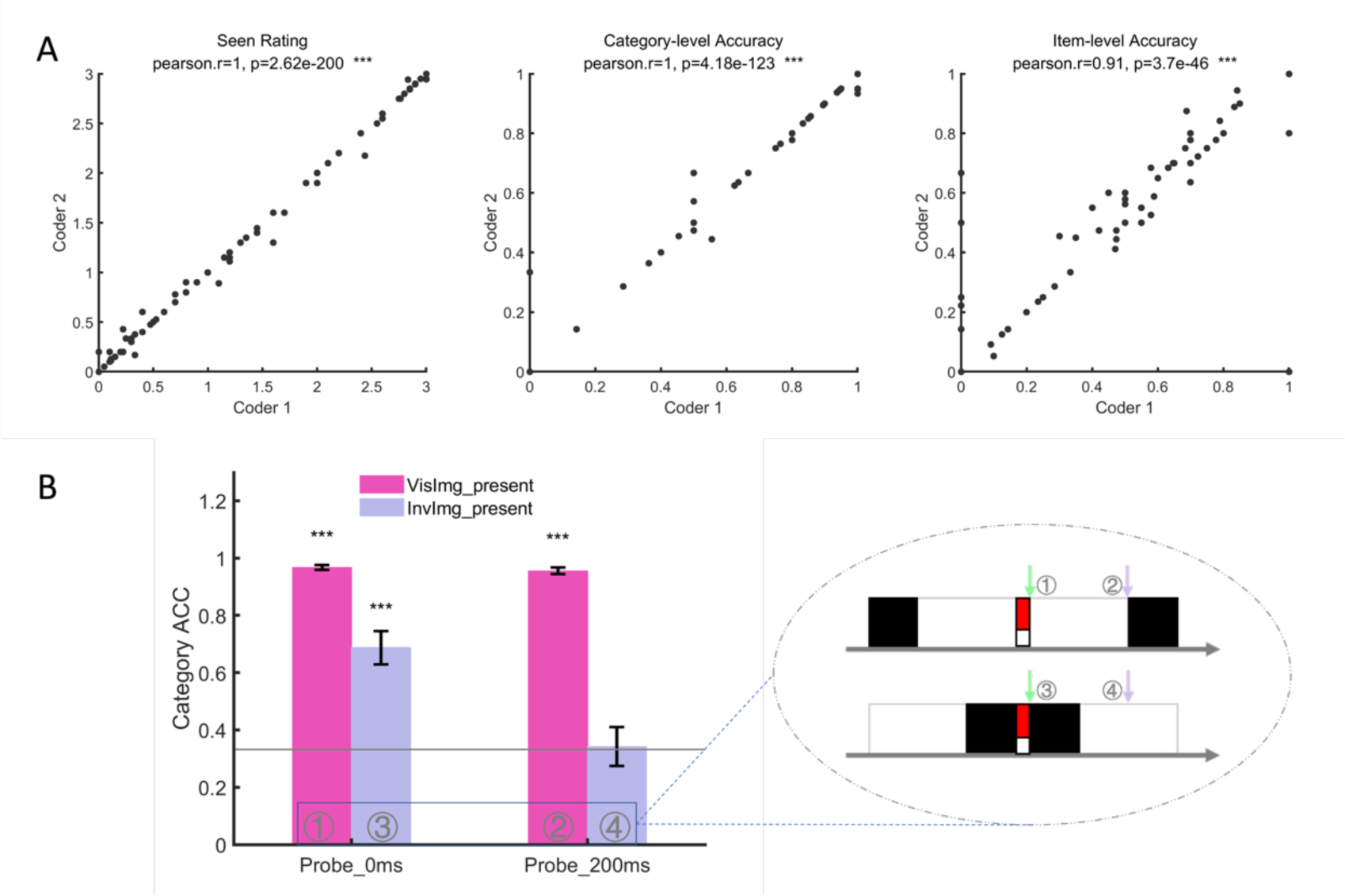
Reliability of coding for oral report and awareness check for probe cues of Exp.1. (A) For each piece of participants’ oral report in probe trials, two independent coders coded visibility rating (left panel), category-level accuracy (middle panel) and item-level accuracy (right panel) with high reliability. For each panel, each dot represents a combined condition of image visibility and probe onset time for a certain participant. For each of the three indices, Pearson correlation analysis revealed near to perfect reliability between two coders. (B) Taking the category-level accuracy averaged across two coders as an objective index of visibility, we found categorization performance varied with combination of visual mask and probe timing (as shown in the suspended ellipse). In condition 1 and 2 where images were weakly masked, optimal accuracy evidenced they were truly visible. By contrast, in condition 4 when images were closely masked by both the forward and the backward masks, the chance-level accuracy suggests that they were completely invisible. In condition 3 when images were masked by forward mask only, the above-chance moderate accuracy suggests there still existed residual awareness for the presented image because the backward mask had not yet been presented. *, p<0.05; **, p<0.01; ***, p<0.001.

**Fig.S2.**
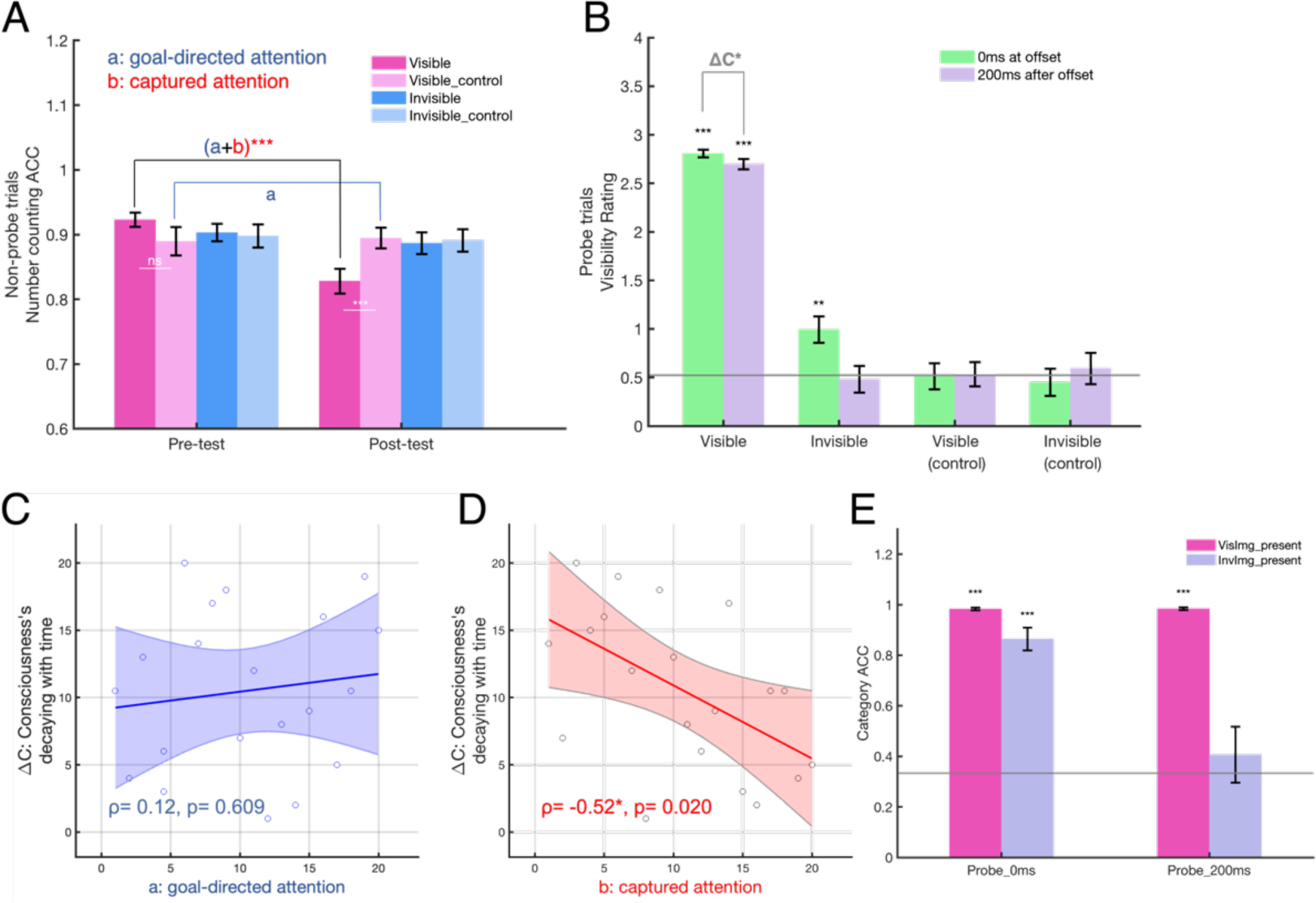
Replication study for main finding of Exp1 that captured attention specifically and proportionally correlates with subjective rating of image visibility. To further test for the link between captured attention and conscious experience, a follow-up experiment was designed and performed using roughly the same paradigm and comparable sample size (20 healthy colleague students, 16 females, age=23.1 ± 1.88 years) but recruiting a different experimenter and two new audio encoders. The major difference lied in the instruction that we provided for participants. That is, we required them to more strictly set priority the counting task and strictly avoid being distracted by peripheral images as best as they can. In extreme cases, it was even acceptable for one to perceptually miss all the images nested in probe trials only if they paid their best attention to counting. Otherwise, if attentional distraction occurred a decrease of counting performance would lead to a decrease of their monetary reward they received at last (note that this ‘punishment’ rule served only as a cover story which did not work in reality). In this way, we aimed to further test the relationship between captured attention and conscious experience in the scenario when the amount of goal-directed attention is largely diminished. We predicted that in this context the amount of captured attention might not necessarily co-diminish with the decreased amount of goal-directed attention. Following results support this view. (A) In the pretest session (i.e., without probe), counting performance did not vary with visibility (F(_3, 57_)= 1.32, p=0.277), confirming that in the pretest session peripheral salient images failed to capture attention. In the posttest session (i.e., with probe), though there was a general decrease (t(_18_)=-2.22, p=0.038) on counting performance in the posttest session relative to the pretest baseline, this general decrease was mainly contributed by visible image trials, whereas performance for visible control, invisible image or invisible control trials exhibited no significant decrease relative to the pretest baseline (uncorrected p>0.1) after we strictly required participants to avoid being distracted. This finding indicates that, expectedly, the amount of goal-directed attention had largely diminished to a minor level (non-detectable). If visible image was shown, however, we replicated the effect of attentional capture even in the near absence of goal-directed attention (also see (*2*)). That is, visible image trials relative to invisible image trials drastically encountered an additional decrease of task performance (interaction: F(_1, 19_)=14.89, p=0.001) which was irrespective of categorical contents (F(_2,38_)=0.44, p=0.650), confirming that when the image became task-relevant and thus attention became available, visibility-related (visible versus invisible) signal automatically and additionally captured attention. Together with Exp.1 reported in the main text, findings here further lead to the conclusion that top-down goal-directed attention and bottom-up attentional capture are two distinct processes. Following prior procedure, we subtracted the amount of attention generally divided to visible control trials (component a: goal-directed attention, t(_18_)=-0.28, p=0.786) from that of visible image trials (component a+b: goal-directed plus captured attention, t(_18_)=5.13, p<0.001) to avoid any confounding from masks, thus obtaining a clean measure of the amount of captured attention (component b). Following results confirm that it is the captured attention that contributes to conscious experience. (B) For 16% probe trials, the decrease of seen ratio (t(_19_)=-2.82, p=0.011) for 200ms versus 0ms probes was observed again, indicating that the consciousness trace was decaying over a 200ms temporal gap. (C and D) Most importantly, we replicated the main finding that the magnitude of consciousness’s decaying over time was negatively correlated with the amount of captured attention (spearman ρ= −0.52, p=0.020, Fig. 1G) but nonsignificant for goal-directed attention (Fig.1F, spearman ρ= 0.12, p=0.609). (E) In line with Fig.S1B, above-chance and chance-level category classification accuracy proved visible and invisible images were indeed consciously and unconsciously perceived, respectively. ns, p>0.05; *, p<0.05; **, p<0.01; ***, p<0.001.

**Fig.S3.**
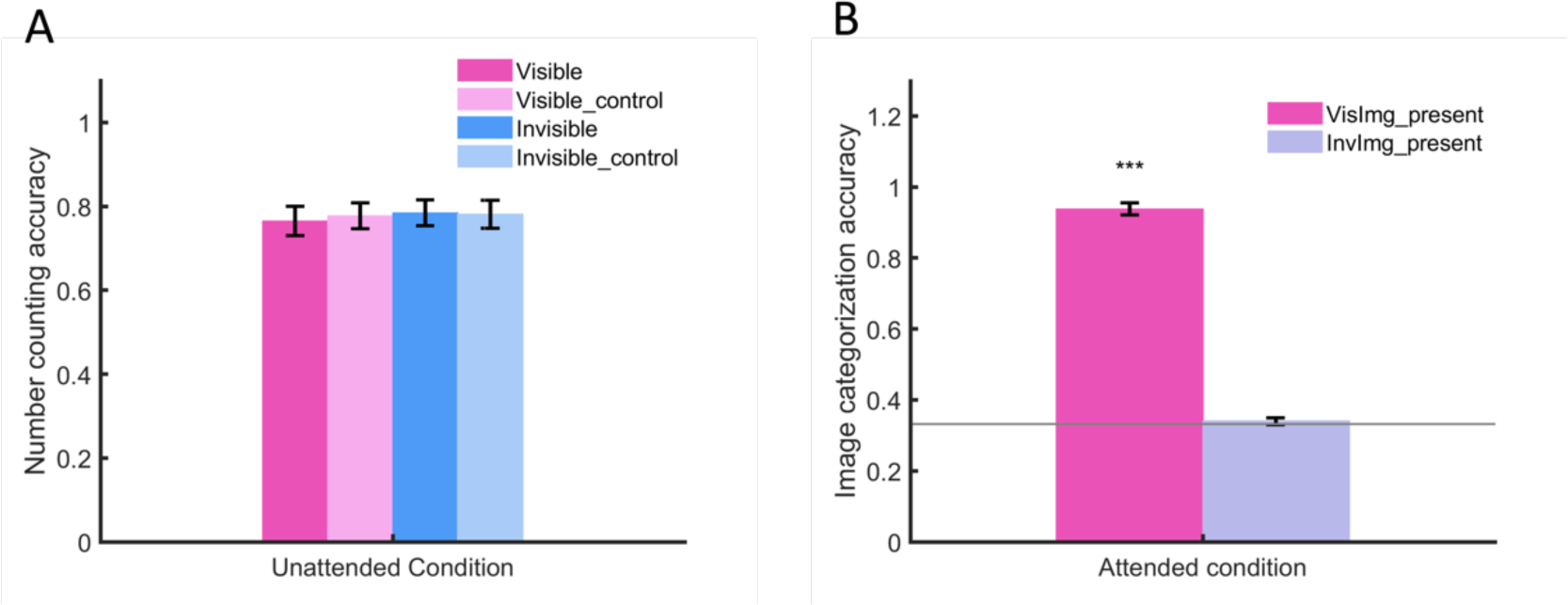
Performance of the number-counting task in unattended condition and image categorization task in attended condition by epilepsy patients of Exp.2. (A) In the unattended condition, counting performance did not vary with image visibility (F(_3, 54_)= 0.95, p=0.424), and performance for visible trials did not differ with that for its matched control (t(_18_)=-1.00, p=0.328). These results collectively suggest that visible signal failed to capture attention in the unattended condition of Exp.2, agreeing well with the unattended pretest session of Exp.1. (B) In the attended condition, patients performed an image categorization task instead. Categorization accuracy was tested against the chance level (1/3) using one-sample t-test, revealing optimal accuracy for visible images (t(_18_)=35.41, p=4.25e-18) and chance-level accuracy for invisible images (t(_18_)=0.99, p=0.335). These results convinced us that visual masking was successful and in the attended condition patients did pay sufficient attention to peripheral image as expected. ***, p<0.001. Error bar represents SEM across patients.

**Fig.S4.**
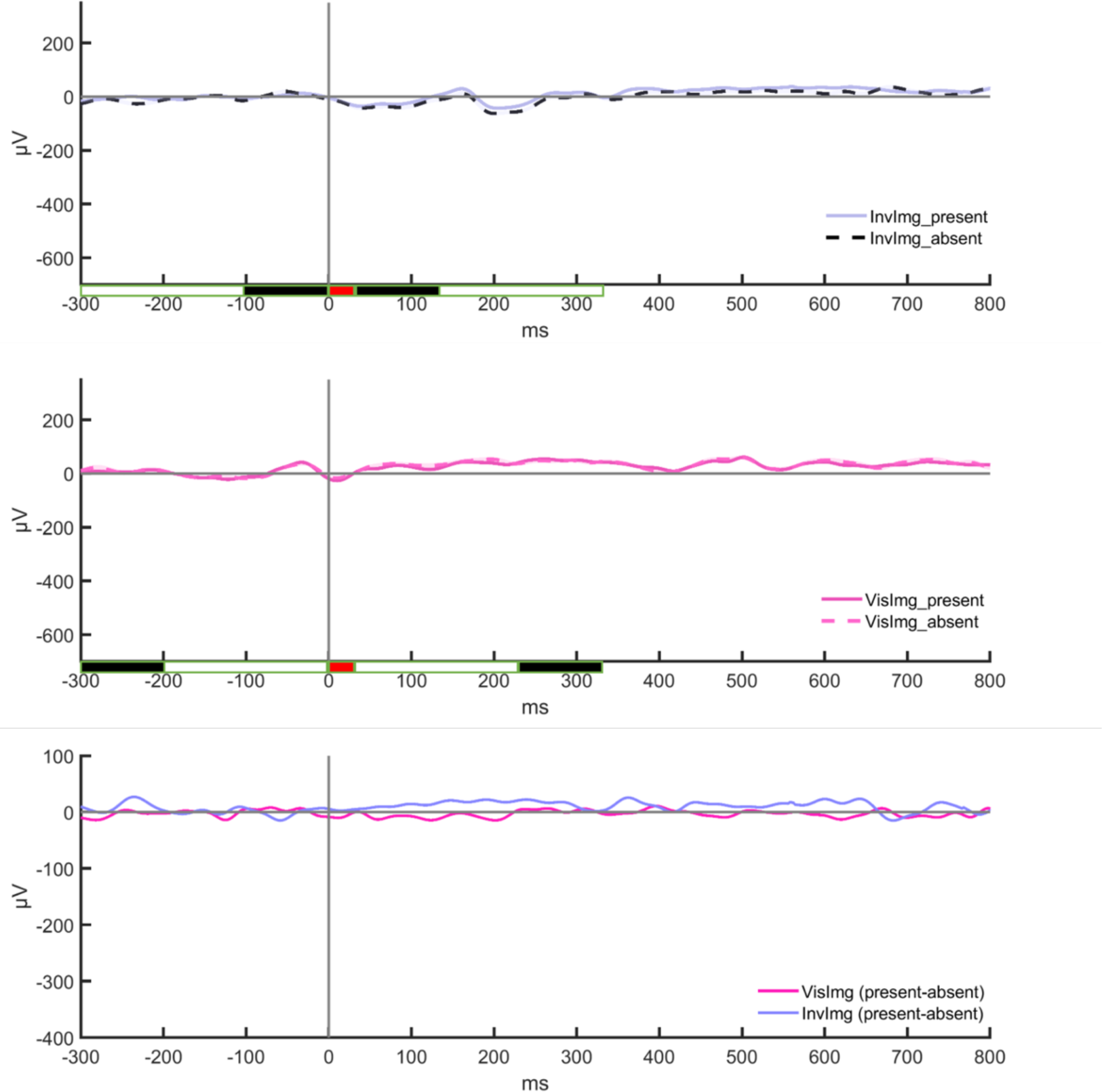
Ipsilateral iERP response of the same representative V1 contact in the unattended condition. The upper and middle rows show V1 response to invisible and visible images, respectively, by evaluating the difference between solid and dotted lines. Tied to the x-axis, rectangles face-colored in white, black and red indicate the presentation of a blank screen, a visual mask and a target image (or its blank control), respectively. Note that the dotted lines indicate background iERP activity evoked by masks only. The bottom row compared V1 response to visible versus invisible images with each subtracting the background iERP. Cluster-based permutation test revealed no clusters showing significant difference between two lines for each panel (corrected p > 0.05). To facilitate comparison between ipsilateral response and contralateral response, limit of the y-axis was kept same with panels of Figure. 2A as shown in the main text. This result suggests that, without attention, V1 at the ipsilateral site could be largely non-responsive to the peripheral visual stream, even if in the ‘visible’ timing structure both masks and image were of strong sensory intensity.

**Figure.S5.**
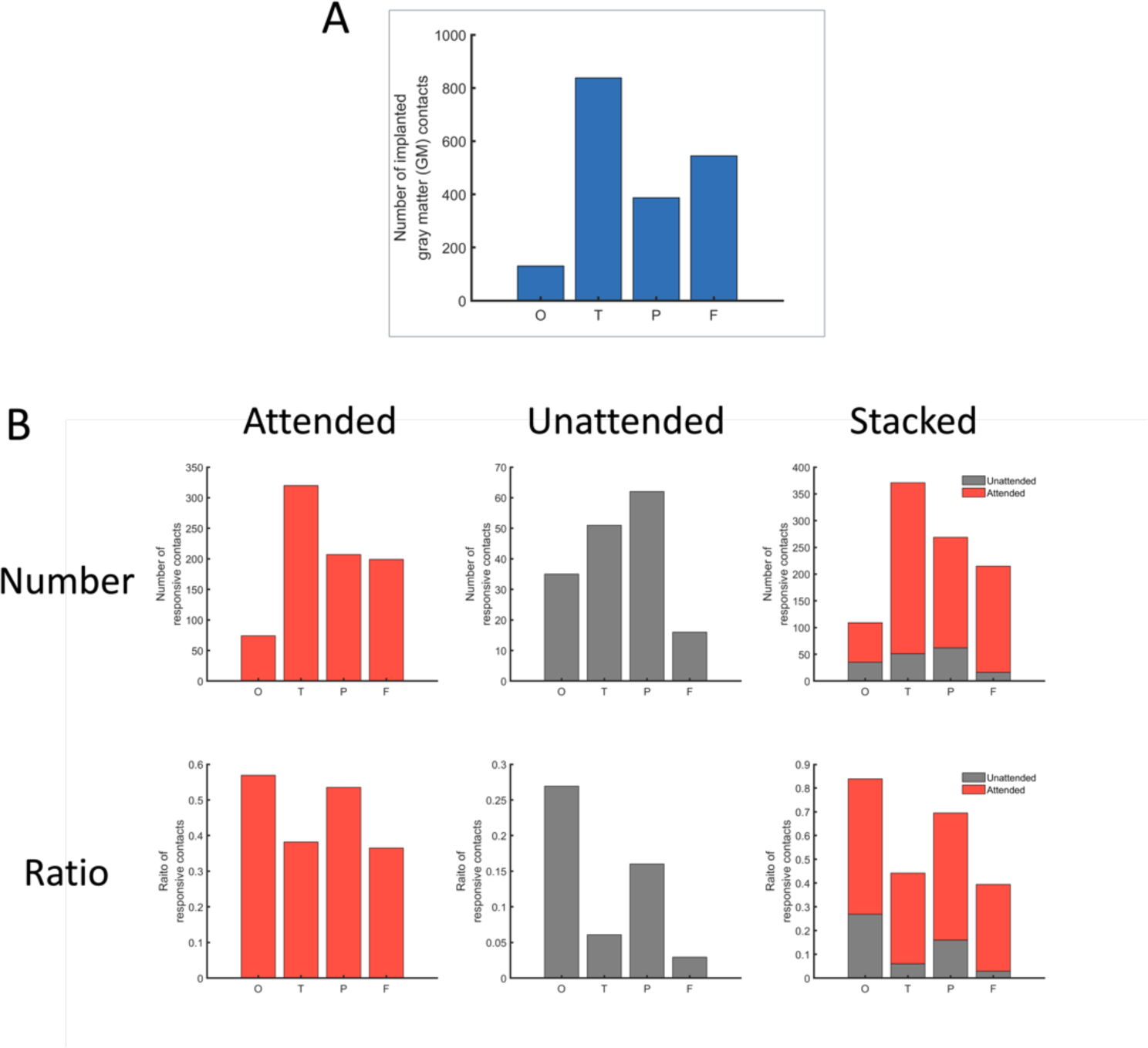
Number (upper row) and ratio (lower row) of responsive contacts located at each of the four lobes. (A) Number of contacts that were successfully implanted to gray matter (GM) tissues. The present study selected responsive contacts from GM only, while contacts that were identified as located in the white matter or out of brain were excluded from further analysis. (B) Distribution of responsive contacts was shown separately for attended (left column) and unattended (middle column) condition, respectively, as well as in their stacked view (right column). In our task designed to tackle the attentional integration process into visual consciousness, the highest ratio of responsive contacts was found at the occipital lobe irrespective of attended or unattended conditions.

**Fig.S6.**
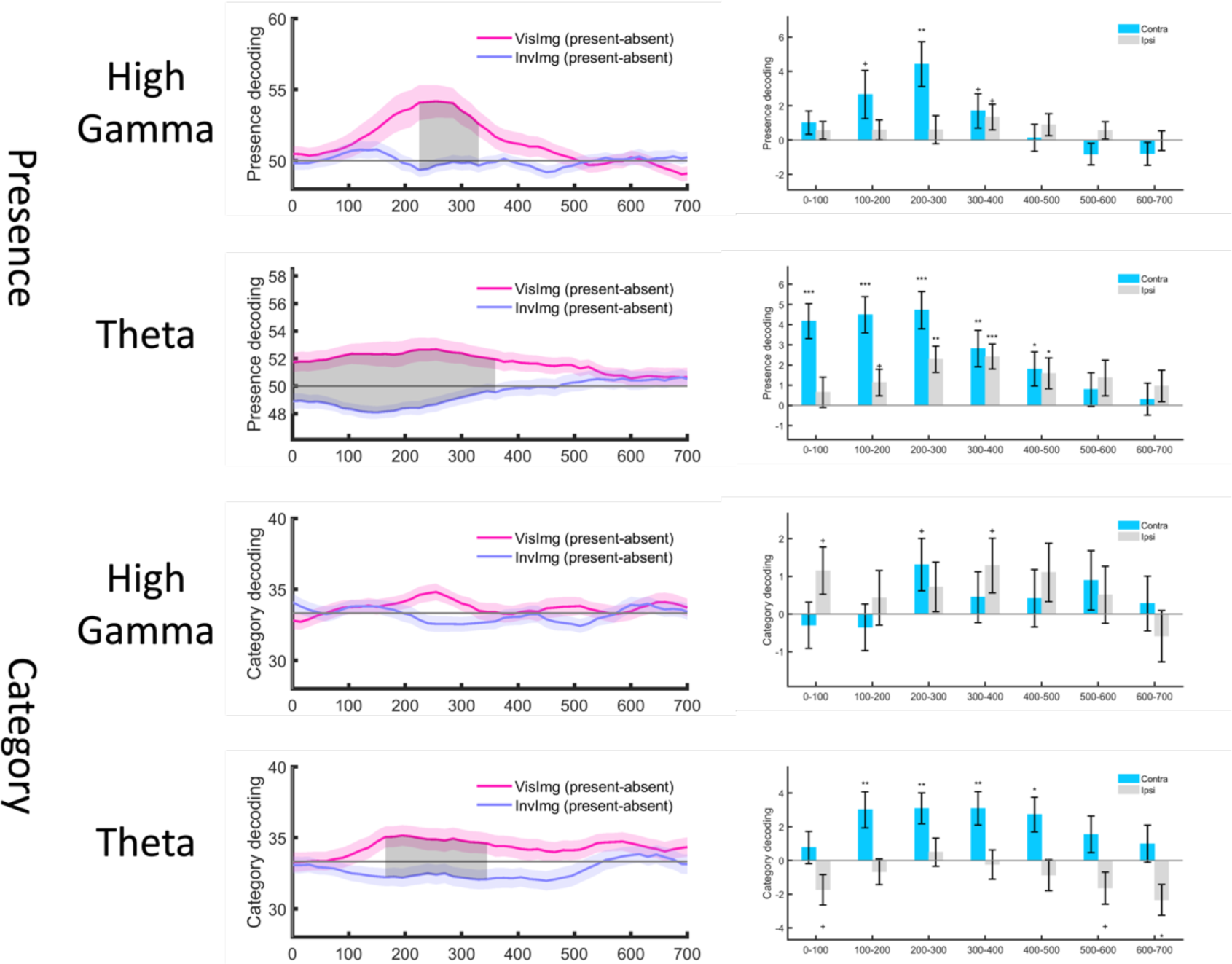
Different contributions of the high-gamma (70-150Hz) and theta (3-5Hz) oscillations at the posterior brain in decoding binary presence (upper blue dotted square) and categorical contents (lower green dotted square) of preconscious signal in the unattended condition. Inside each square, panel figures in the first column showed decoding performance for both visible (light red lines) versus invisible (light blue lines) signals, with each subtracting the performance of image-absent trials from that of image-present trials (i.e., present-absent). The gray shaded areas indicated time clusters of significant difference between visible versus invisible signals (corrected for multiple comparisons using the cluster-based permutation test, p<0.05), i.e., preconscious signal observed in the unattended condition. For this signal, panel figures in the second column further showed differential decoding performance (visible - invisible) of both the contralateral and ipsilateral hemispheres of the posterior brain in a bin-wise manner. Decoding performance was averaged across each bin of 100ms bin width and step size. Statistical significance of each bar value differing from 0 was indicated by an asterisk shown tied to the error bar (one-sample t-test, uncorrected). +, p<0.1; *, p<0.05; **, p<0.01; ***, p<0.001; otherwise, p>0.1. Error bar indicates SEM across responsive units of the posterior brain in the unattended condition. As shown, when attention was not engaged, responsive units at the posterior brain can reliably decode binary presence of preconscious signals by using both theta and high gamma oscillations, particularly by contralateral hemisphere of the posterior brain. Similarly, fine-grained categorical contents of the preconscious signal were also preferentially decoded by the contralateral hemisphere. The difference was that it was the slow theta oscillation rather than the high-gamma oscillation that more pronouncedly contributed to decoding these fine-grained contents (although a trend effect was observed for contralateral hemisphere high gamma oscillations at the 200-300ms bin).

**Fig.S7.**
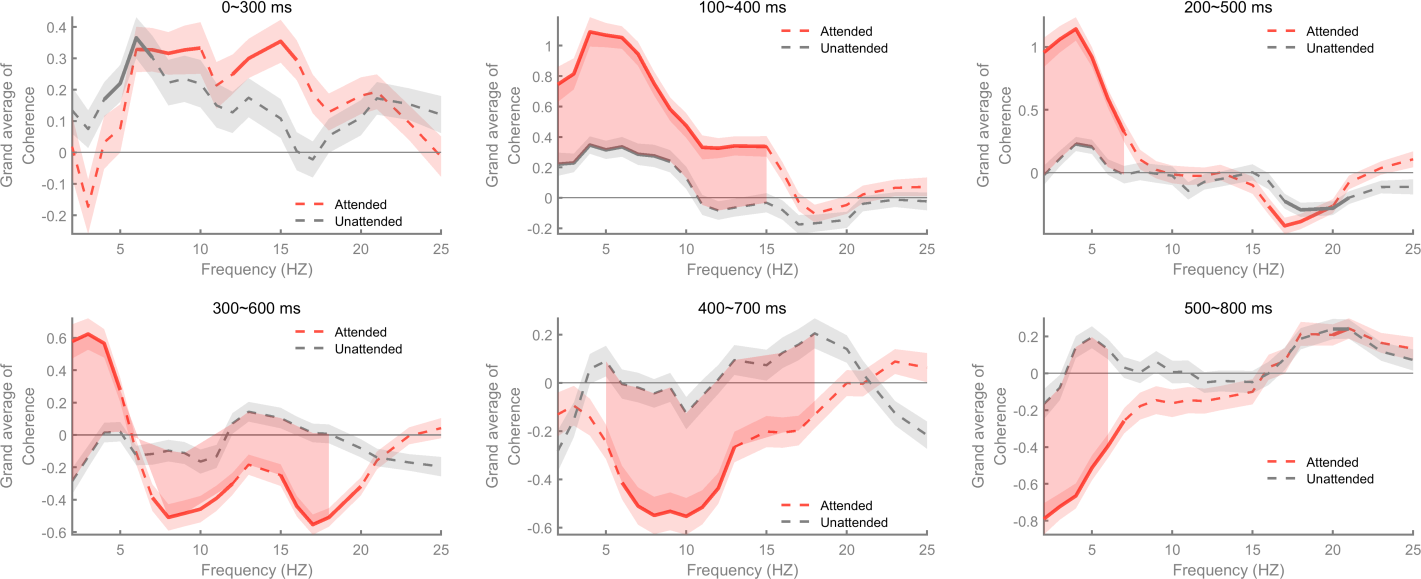
The grand averaged coherence value which measures the difference between visible versus invisible trials (visible-invisible) varied as a function of frequency and time bins in both attended (red) and unattended (gray) conditions. Note that this measure was calculated as adjusted imaginary coherence which may produce values exceeding 1 (See Method for more details). For each condition and each bin of 300ms window size, we averaged consciousness related coherence values across all lobe pairs while correcting for data homogeneity between contacts from the same electrode (see Methods). Solid lines indicate significant change in attended and unattended conditions, respectively (cluster-based permutation test, p<0.05). Dotted lines indicate non-significant results (cluster-based permutation test, p>0.05). Shaded area indicates significant difference between the attended and unattended conditions (cluster-based permutation test, p<0.05). As shown in this figure, attentional modulation on visibility-related (visible-invisible) phase coherence was mainly observed in 4-Hz peaked theta band (both unattended and attended conditions) in earlier bins and 10-Hz peaked alpha band (mainly in attended condition) in later bins.

**Fig.S8.**
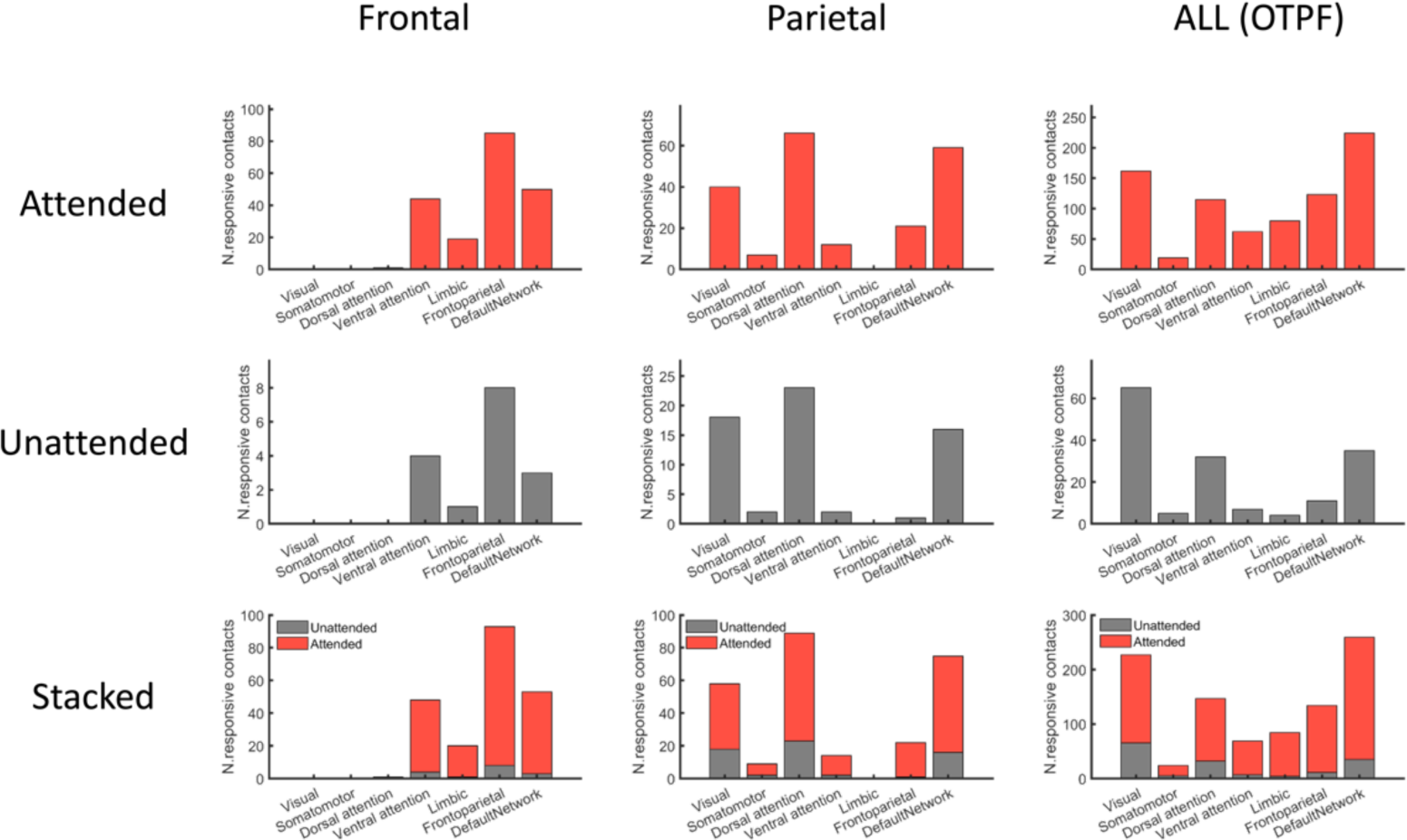
Distribution of responsive contacts in both attended (top row) and unattended (middle row) conditions coming from seven large-scale cerebral networks (Yeo-7 template (*79*)). Stacked bars (bottom row) showed how much the number of responsive contacts increased due to engagement of attention in each of the network. This relabeling process was conducted for responsive contacts of the frontal lobe (left column) and parietal lobe (middle column) of special interest, as well as for all responsive contacts in human neocortex (right column). As shown in this figure, for both attended and unattended conditions, responsive contacts from the frontal lobe (left column) were mainly from the frontal-parietal network, while the parietal lobe ones (middle column) were mainly from the dorsal attention network. Counting all responsive contacts together (right column), we found that the network containing the largest number of responsive contacts changed from the visual network (unattended condition) to default network (attended condition) after attention was engaged.

**Fig.S9.**
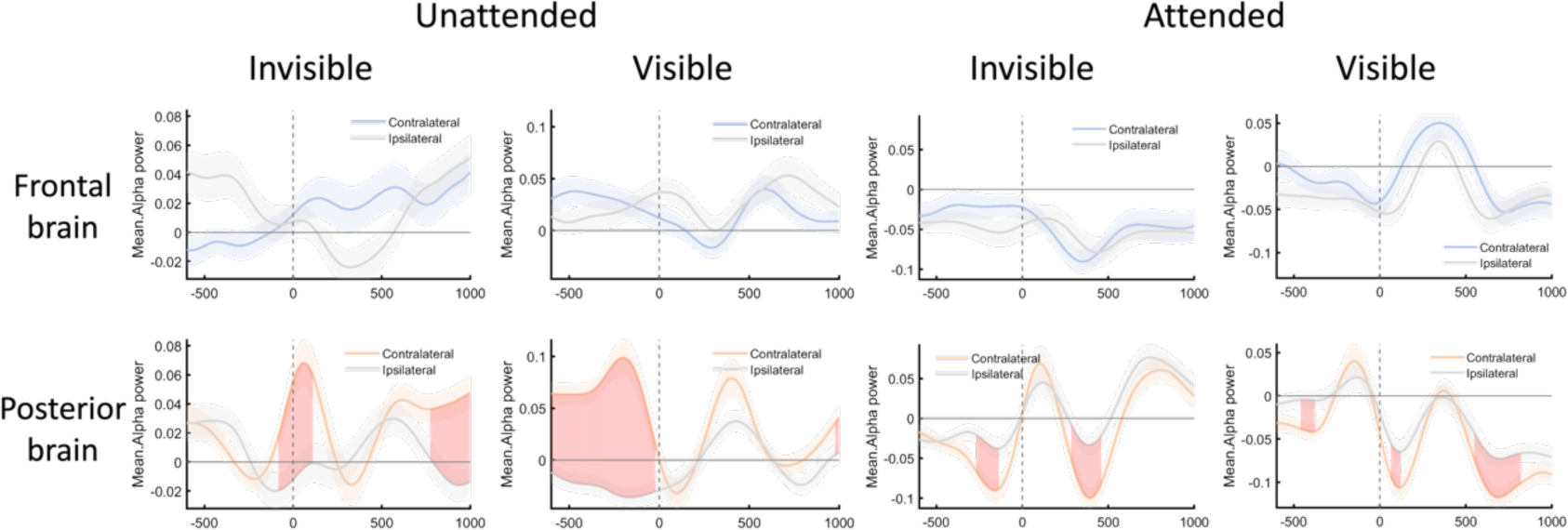
Attention-induced pre-stimulus lateralized distribution (contralateral versus ipsilateral) of normalized alpha power for each of four combined conditions across visibility and attention, for each of the frontal brain (top row) and the posterior brain (bottom row). Note that visible and invisible images were nested into different timing structure, which might have changed pre-stimulus alpha activity. To evaluate inter-trial variance of alpha activity, power values were first normalized across all trials before being sub-grouped into different conditions to evaluate their variance during the pre-stimulus period. For this aim, baseline correction procedure was not performed to preserve baseline activity. Since pre-stimulus visual stream was the same between image present trials and image absent trials (i.e., mask only trials), the normalized alpha power value of these two types of trials were averaged for each of the eight combined conditions (corresponding to eight panels). For each panel, two lines displayed represent contralateral and ipsilateral response, respectively. Shaded area around each line indicates SEM across responsive units (see Method), and the shaded area inter-between two lines indicates significant difference between them caused by neural lateralization (cluster-based permutation test, p<0.05). Zero point in the x-axis indicates image onset. Notably, the location of the peripheral stream was the same across trials of a block (randomly determined between blocks). In the unattended condition, this setting facilitated patients to deliberately neglect the peripheral stream so as to avoid attentional distractions. In line with this view, we observed that the power of inhibitory alpha oscillation was concentrating to the contralateral side of the posterior brain, to which signals of the peripheral visual stream would be initially imported. This may hinder these incoming sensory signals to capture attention. In the attended condition, however, participants were required to direct covert attention to peripheral streams. This might render the posterior brain into a preparation state. In line with this view, we observed a general decrease of alpha power at the posterior brain after attention was engaged (pink line of bottom panels), especially for the contralateral hemisphere. Alpha power there became even lower than the ipsilateral hemisphere, rendered into an opposite pattern compared with that of the unattended condition. These results suggest that the contralateral hemisphere was suppressed in the unattended condition and facilitated at the attended condition. In sharp contrast with the posterior brain, results above revealed no clear evidence favoring alpha power lateralization at the frontal brain in any combined conditions of visibility and attention.

**Fig.S10.**
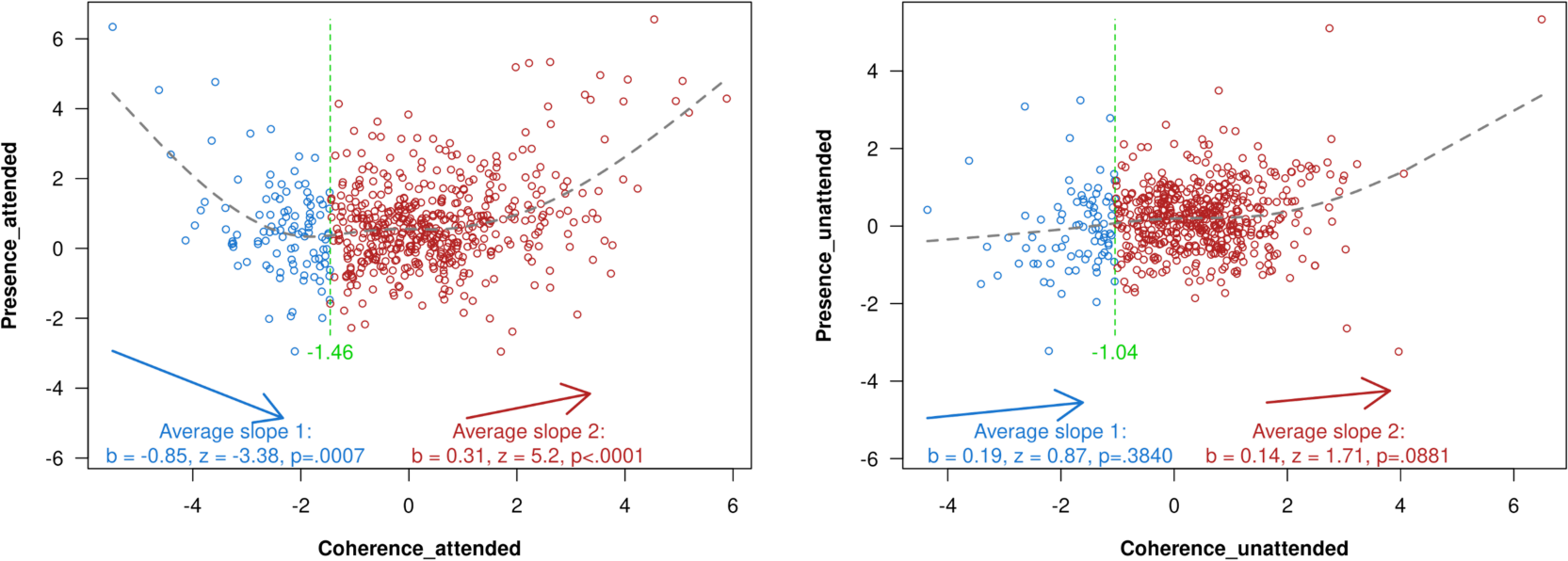
A more rigorous way testing for the U-shape relationship displayed in Figure.3E of the main text. In Figure.3E, we investigated a U-shape relationship between the predictor (referred to as x) and the dependent variable (referred to as y), where x was a vector measuring visibility-related (visible-invisible) change of theta coherence values for each pair of responsive unit and y was a vector of the same length measuring to what extent theta coherence values could discriminate between visible versus invisible trials (training SVM models, see Methods for more information). Both x and y were calculated using data within the early window (0-300ms) at the posterior brain. Methodologically, this U-shape line was drawn based on a simulation fitting a quadratic polynomial curve to the data which was completed using the fit.m function in MATLAB with a fittype of ‘poly2’. This function takes the form f(x) = p1*x^2 + p2*x + p3, where p1, p2, and p3 are the coefficients of the polynomial. The fit.m function finds the best-fitting polynomial curve to the data points, minimizing the sum of the squares of the differences between the data points and the curve. Though this approach testing U-shaped functions via quadratic regressions is widely used, it imposes an arbitrary functional-form assumption that in some scenarios can lead to a 100% rate of false positives (e.g., the incorrect conclusion that y = log(x) is U shaped). Here, as a more rigorous means to test for this U-shape relationship without a functional-form assumption, the two-lines test (*85*) was used to further test this U-shaped relationship. Basically, this approach estimates two regression lines, one for low and one for high values of x. A breakpoint xc was first set using a “Robin Hood” algorithm which seeks to obtain higher power to detect a u-shape if it is present. More details and an application procedure for running the two-lines test are available at http://webstimate.org/twolines. After setting a breakpoint, the two-lines test runs two interrupted regressions, one which includes the breakpoint in the first segment, then one which includes it in the second. Each regression results in a parameter slope b, as well as another two Z and p values evaluating its statistical significance. If the resulting two slopes have opposite sign, and are individually statistically significant, the test rejects the null hypothesis that there is no u-shaped (nor inverted u-shaped) effect. As a result, in the attended condition (left panel), we found that the first (blue) line had negative slope while the second (red) line had positive slope, and both slopes were statistically significant (p<0.001). By contrast, in the unattended condition (right panel), the slope of the first and the second lines were both positive, and neither of them reached statistical significance (p>0.05). Collectively, the U-shape relationship indeed existed but only existed in the attended condition. This finding implies that attention initiates a functional reorganization process at the posterior brain, rendering functionally coupled (positive coherence) and decoupled pairs (negative coherence) simultaneously detecting the presence of conscious signals at the very beginning.

**Fig. S11.**
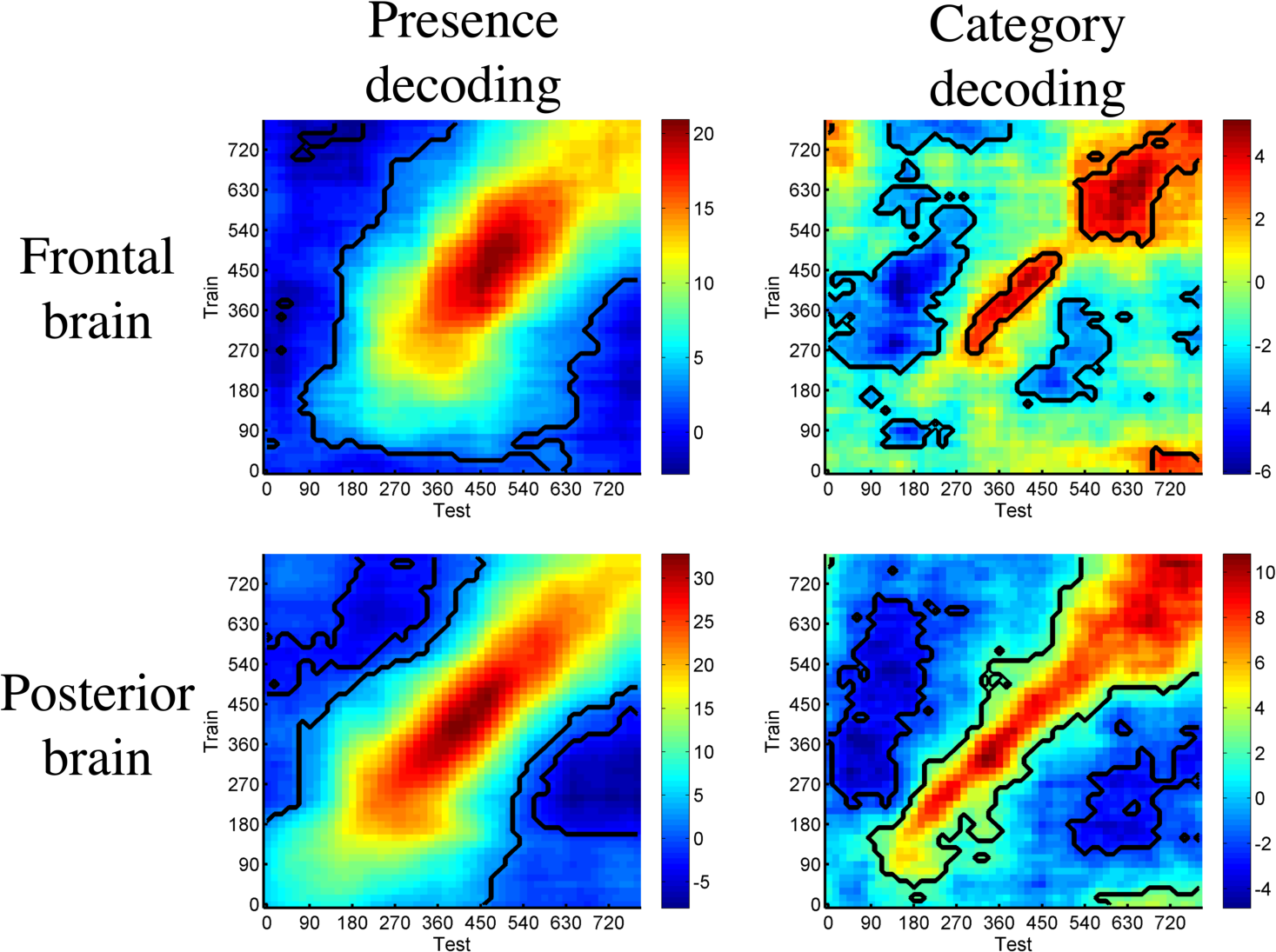
Temporal generalization of neural representations of consciousness in the attended condition. Generalization analysis was performed for both the frontal brain (upper row) and posterior brain (lower row) with respect to coarse-grained decoding of consciousness presence (left column) and fine-grained decoding of categorical contents (right column). Contacts used in this figure were the ones that were responsive in the attended condition. For each panel and time bin, SVM models were trained first. Then, the trained models were used to test data from all bins, producing a train time * test time matrix evaluating to what extent neural representations generalize across different time bins. The warm-colored clusters with black contours indicate bin pairs showing significant temporal generalization effect (correcting for multiple comparisons using the FDR method, corrected p < 0.05). As shown, both coarse-grained and fine-grained representations of consciousness appeared earlier in the posterior brain than the frontal brain. After that, coarse-grained representations kept evolving continuously across time (diagonal direction) in both the frontal and posterior brain, and got maintained more durably across time (horizontal and vertical directions) compared with the profile of fine-grained representations. Especially, in terms of fine-grained consciousness contents, there exhibited a sharp discrepancy between the frontal brain and posterior brain. Whereas the posterior brain held early-starting and continuously evolving representations of consciousness contents, the frontal brain started late and showed two temporally discrete clusters whose representations could not generalize to each other. These results further suggest that the frontal brain had acted two intrinsically different responses while representing consciousness contents. As discussed in the main text, we hypothesized that the first and second component of frontal content representations may support reflexive and deliberate action, respectively.

**Fig.S12.**
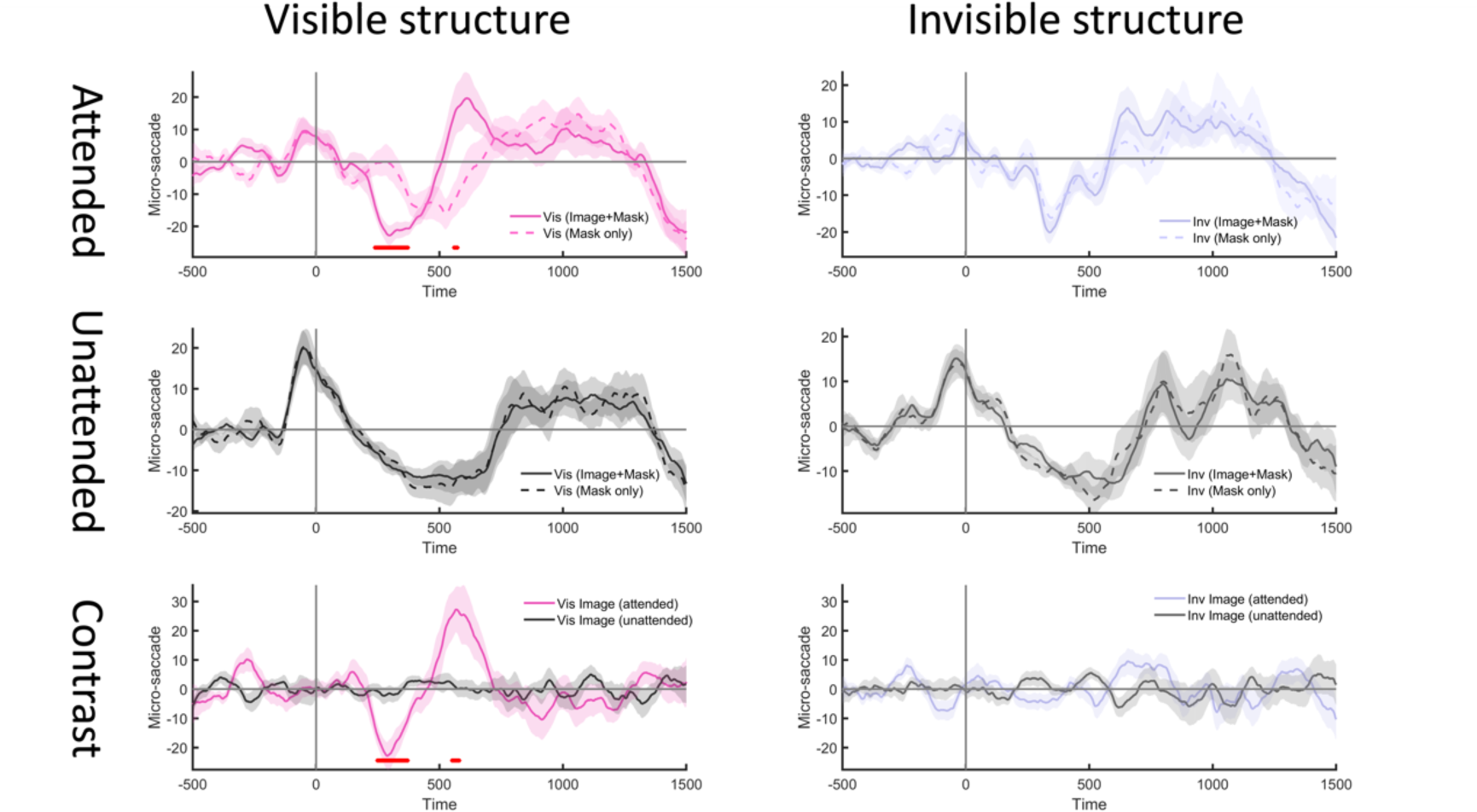
Micro-saccade behavior captured the presence of consciousness signal at the attended rather than unattended condition. Visible and invisible trials had different timing structure. For visible structure (left column) the target image was visible due to weak visual masking, and for invisible structure (right column) invisible due to heavy visual masking. Response to visual masks was measured by mask-only trials. For each structure, image was either attended (first row) or unattended (second row). The contrast between image present trials and mask only control trials was plotted for each attended and unattended condition (third row), to illustrate the effect of attention on micro-saccade behavior in response to image onset. Basically, micro-saccade ratio was first calculated as percentage of trials identified with micro-saccade event for each timepoint. Then, mean percentage of pre-stimulus baseline was subtracted from all other timepoints to evaluate change of micro-saccade ratio caused by target image. Shaded area indicates SEM across subjects after baseline subtraction. Red asterisk indicates timepoint of significant difference between two lines (corrected p < 0.05) of each panel while correcting for multiple comparison tests using cluster-based permutation test. As shown above, micro-saccade behavior captured the presence of consciousness signal at the attended condition only, with micro-saccade ratio first decreasing and then increasing in response to the presentation of visible image. This temporally reversed pattern of micro-saccade response was not observed in the unattended condition.

**Fig.S13.**
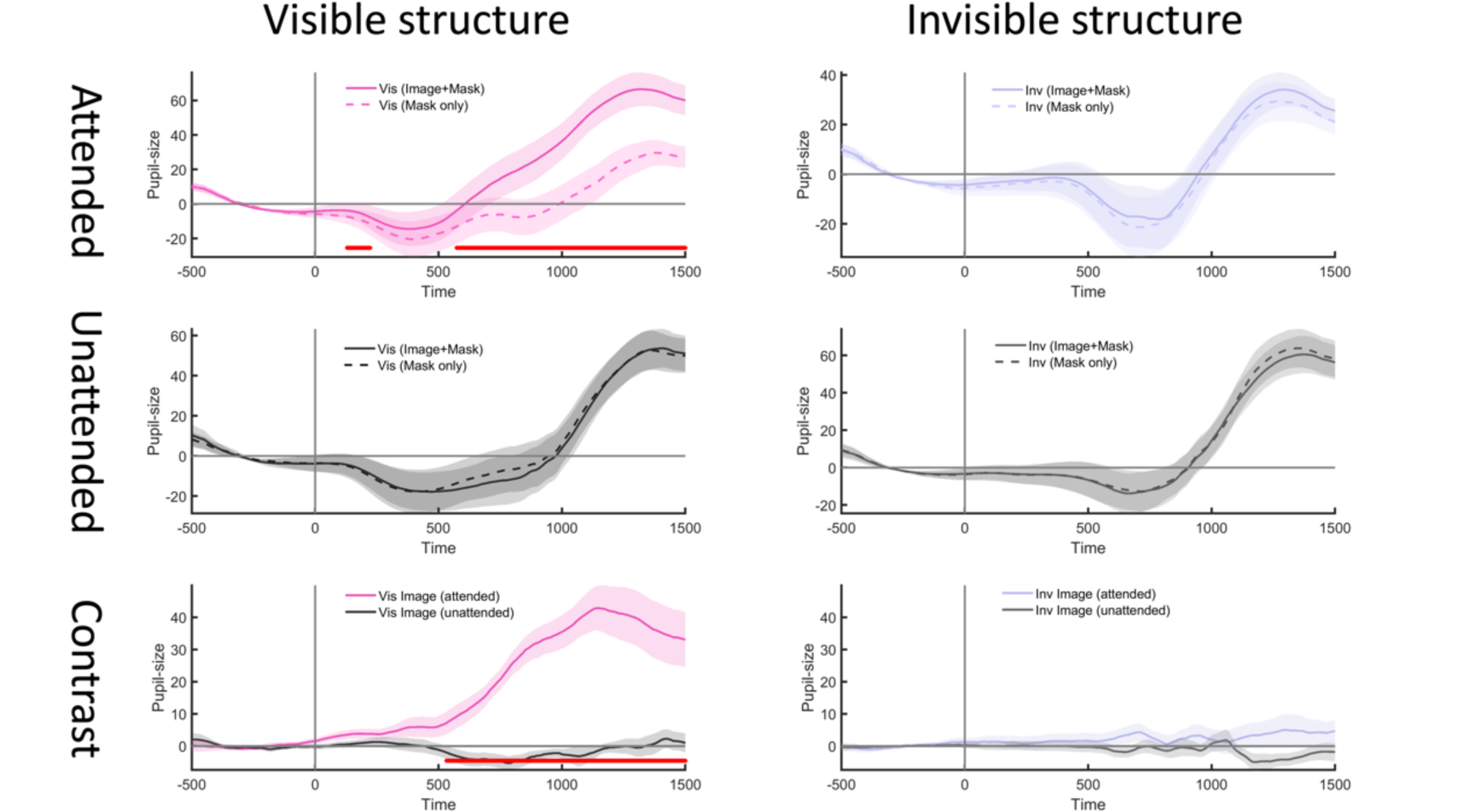
Pupil-size dilated in response to the presence of consciousness signal, an effect only observed at the attended condition too. Annotations here are same as Figure.S11 except for the analysis now was conducted on pupil size (more specifically the diameter of the recorded eye). Shaded area indicates SEM across subjects after baseline subtraction. Red line at the bottom indicates timepoint of significant difference between two lines (corrected p < 0.05) of each panel while correcting for multiple comparison tests using cluster-based permutation test.

**Fig.S14.**
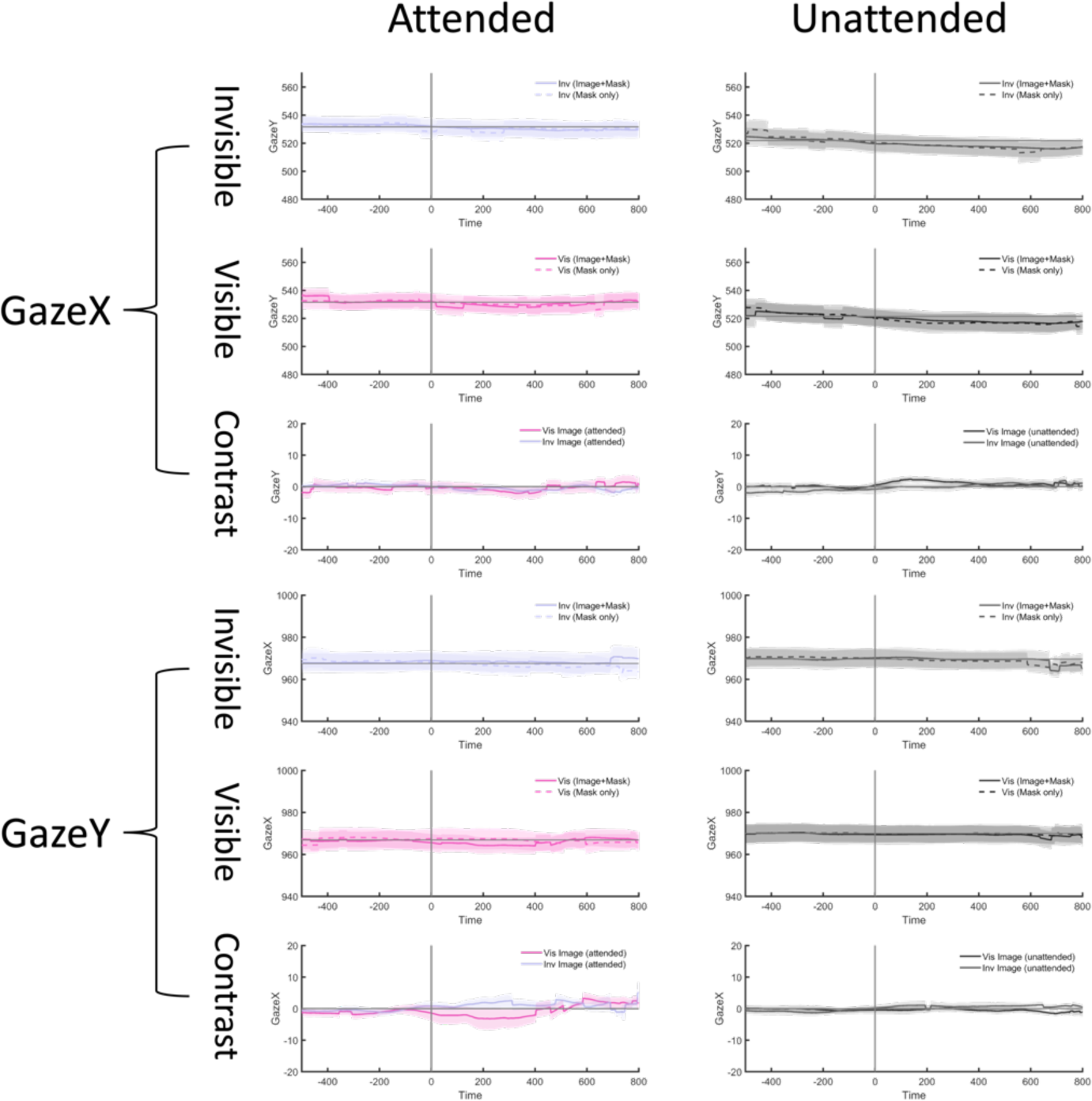
Findings of the present study were not contaminated by difference in pre-stimulus eye gaze location in either horizontal (GazeX) or vertical (GazeY) direction. For each GazeX and GazeY, zero point in the X-axis of each panel figure denotes the onset of target image. In each combined condition of image visibility (first two rows) and attention (two columns), gaze location was equivalent between image trials where target image was presented while surrounded by images and control trials where only masks were presented in the absence of the target image. To detect any difference during the pre-stimulus period, baseline correction was not conducted for this analysis on raw gaze location. As a result, no timepoint passed a liberal threshold of uncorrected p<0.05. As shown in the third row (‘Contrast’), there was no significant difference between the visible and invisible condition after each subtracting mean gaze location values of corresponding control trials (uncorrected p>0.05), indicating comparable eye gaze location between the visible and invisible state of image processing. Thus, the neural dynamics underlying attentional integration into consciousness, as we reported in the present study, were not contaminated by any systematic difference in pre-stimulus eye gaze location.

**Fig.S15.**
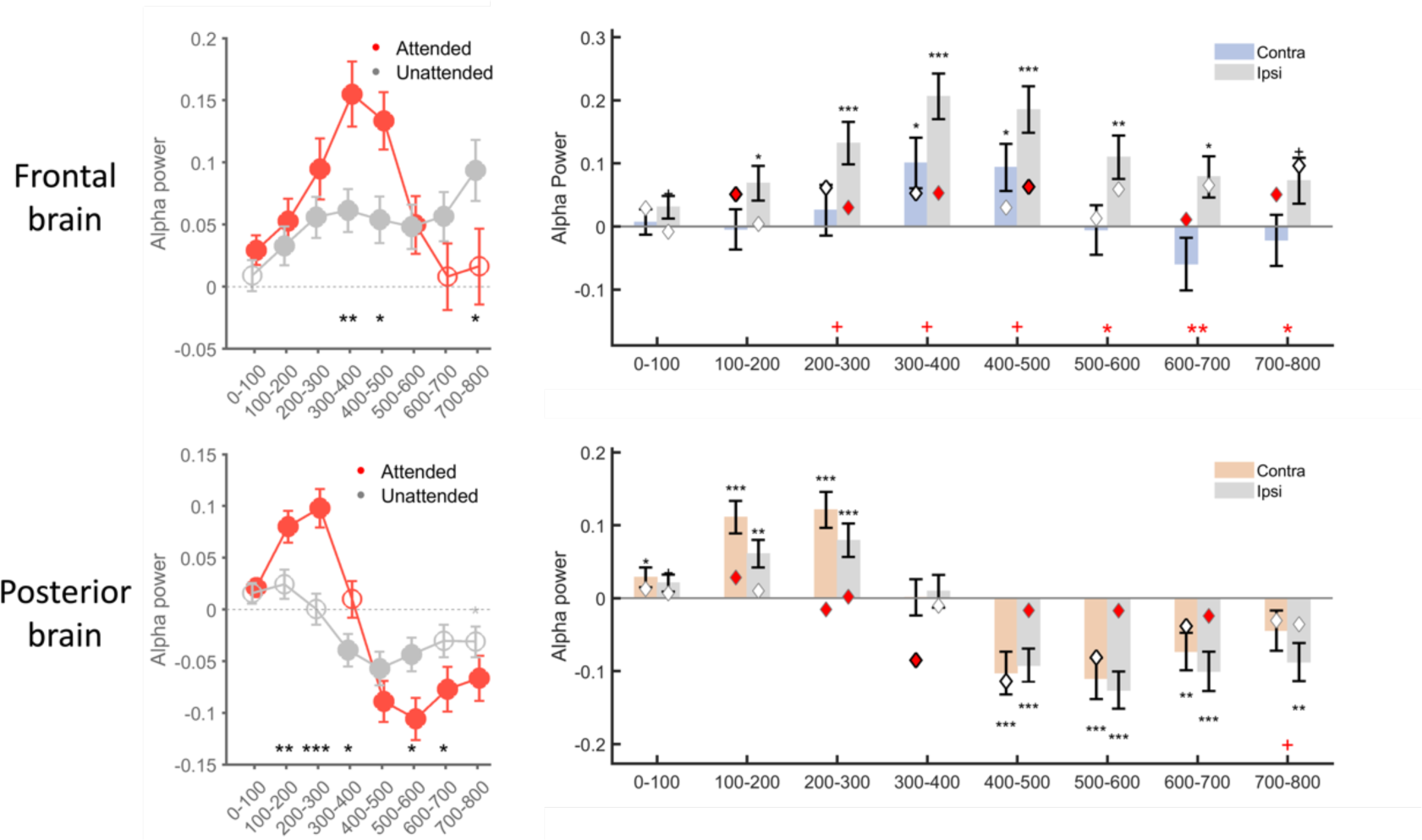
Post-stimulus alpha power distribution in the frontal brain (upper row) and the posterior brain (bottom row). In the left column, visibility-related (visible-invisible) alpha power change was preferentially compared between attended (red) and unattended (gray) conditions. For each condition and each bin, unfilled circles without asterisks indicate non-significant response (t-test, uncorrected, p>0.05), unfilled circles with asterisks indicate significant response without correcting for multiple comparisons (t-test, uncorrected, p<0.05), and filled circles indicate a significant response after correcting for multiple comparisons (t-test, FDR corrected, p<0.05). For each bin, the statistical significance of the contrast between attended and unattended conditions was marked by asterisks (paired t-test, FDR corrected, p<0.05) shown at the middle-bottom line. *, p<0.05; **, p<0.01; ***, p<0.001. Error bar indicates SEM across responsive units. In the right column, alpha power was further compared between contralateral versus ipsilateral hemispheres. Alpha-band power for contralateral versus ipsilateral stimulus was averaged across each bin of 100ms. For each panel, mean alpha power for contralateral versus ipsilateral stimulus was compared, with red symbols shown in the middle bottom indicating statistical difference between two bars in the attended condition (paired t-test, uncorrected, p<0.05). For the attended condition, statistical significance different from 0 was indicated by an asterisk shown above the error bar (t-test, uncorrected, p<0.05). For the unattended condition, mean decoding performance was indicated by the position of the diamond symbol and result of the same t-test against 0 was coded by the diamond symbol (t-test, uncorrected, p<0.05). Diamond with bold and black edges indicates a significant change (uncorrected p<0.05) in the unattended condition while the one with thin and gray edges indicates a non-significant result (uncorrected p>0.05). Beyond that, face color of the diamond indicates the statistical difference between attended and unattended condition, with red face color indicating a significant difference between them (paired t-test, uncorrected p <0.05) while the white face color indicating a non-significant modulation effect (paired t-test, uncorrected p>0.05). To better evaluate any trend effect, correction for multiple comparisons were not conducted. +, p<0.1; *, p<0.05; **, p<0.01; ***, p<0.001; otherwise, p>0.1. Error bar indicates SEM across responsive units. This figure highlights that engagement of attention into consciousness produces different profiles of alpha power activity. In the attended condition, especially, alpha activity differently acts in the frontal brain versus the posterior brain. In the frontal brain, alpha activity is persistently lateralized to the ipsilateral hemisphere, suggesting the ipsilateral frontal brain was inhibited during the whole process. In the posterior brain, however, the two-stage reversed alpha activity which is also comparable between two hemispheres suggests that both hemispheres of the posterior brain encounter an early inhibition followed by a late excitation.

**Fig.S16.**
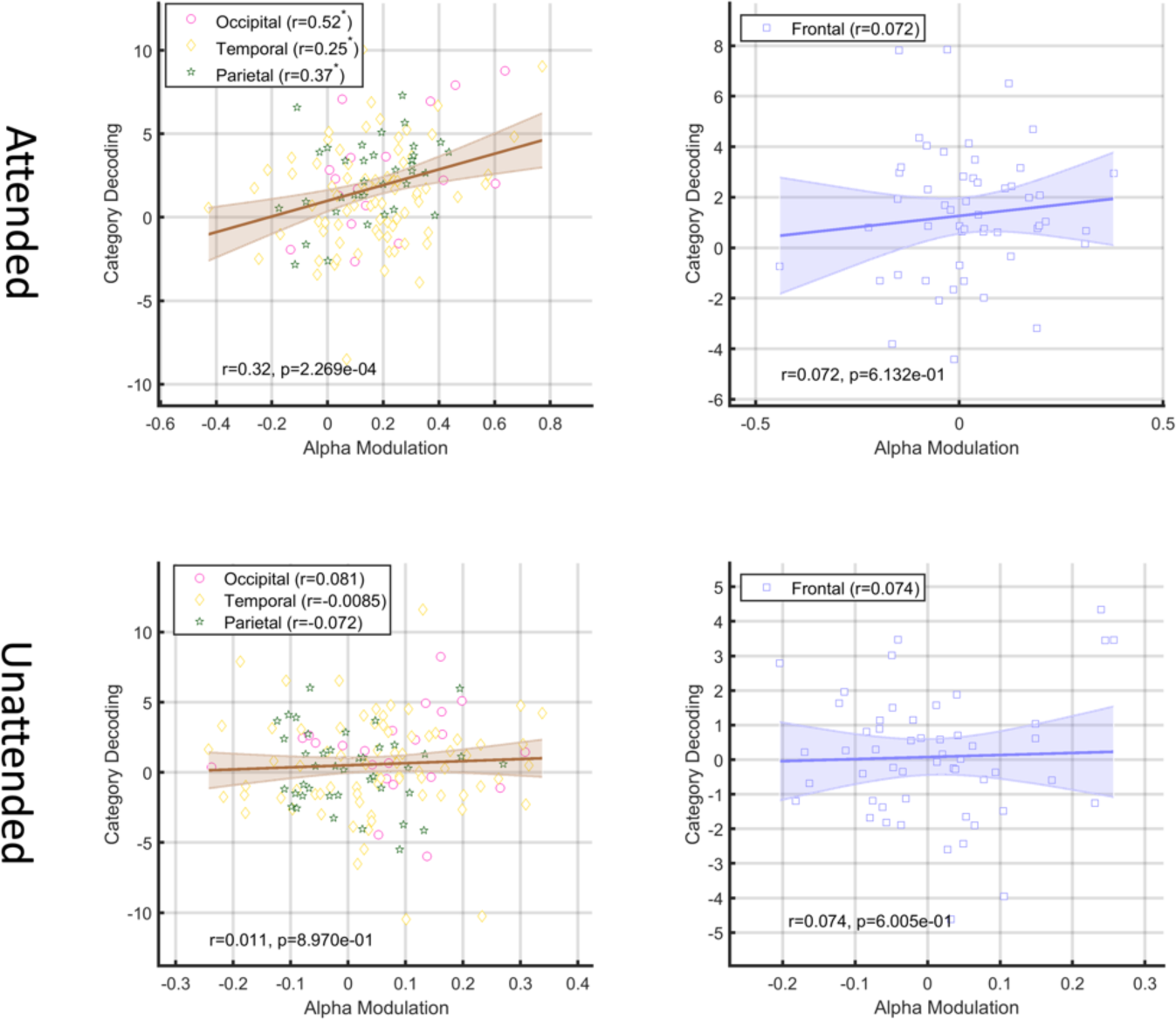
Functional link between visibility-related (visible-invisible) alpha activity and neural representations of consciousness contents. For each panel, alpha modulation effect was calculated as the amplitude of alpha power change by subtracting the mean power of the late stage (400-700ms) from that of the early stage (0-300ms, see Fig.3I and 3K). Then, Pearson correlation was conducted between the amplitude of the alpha modulation effect and the content decoding performance for the posterior brain (left column) and the frontal brain (right column), in both the attended condition (upper row) and the unattended condition (lower row). Same analysis was also conducted for each of the four individual neocortex lobes, with the correlation coefficient (r) delineated within brackets shown at the legend position of each corresponding panel. *, p<0.05. As shown, the two-stage alpha modulation effect is correlated to content decoding performance for the posterior brain, and only significant in the attended condition, which suggests that attention may engage two-stage alpha activity to filter consciousness contents at the posterior brain.

**Fig.S17.**
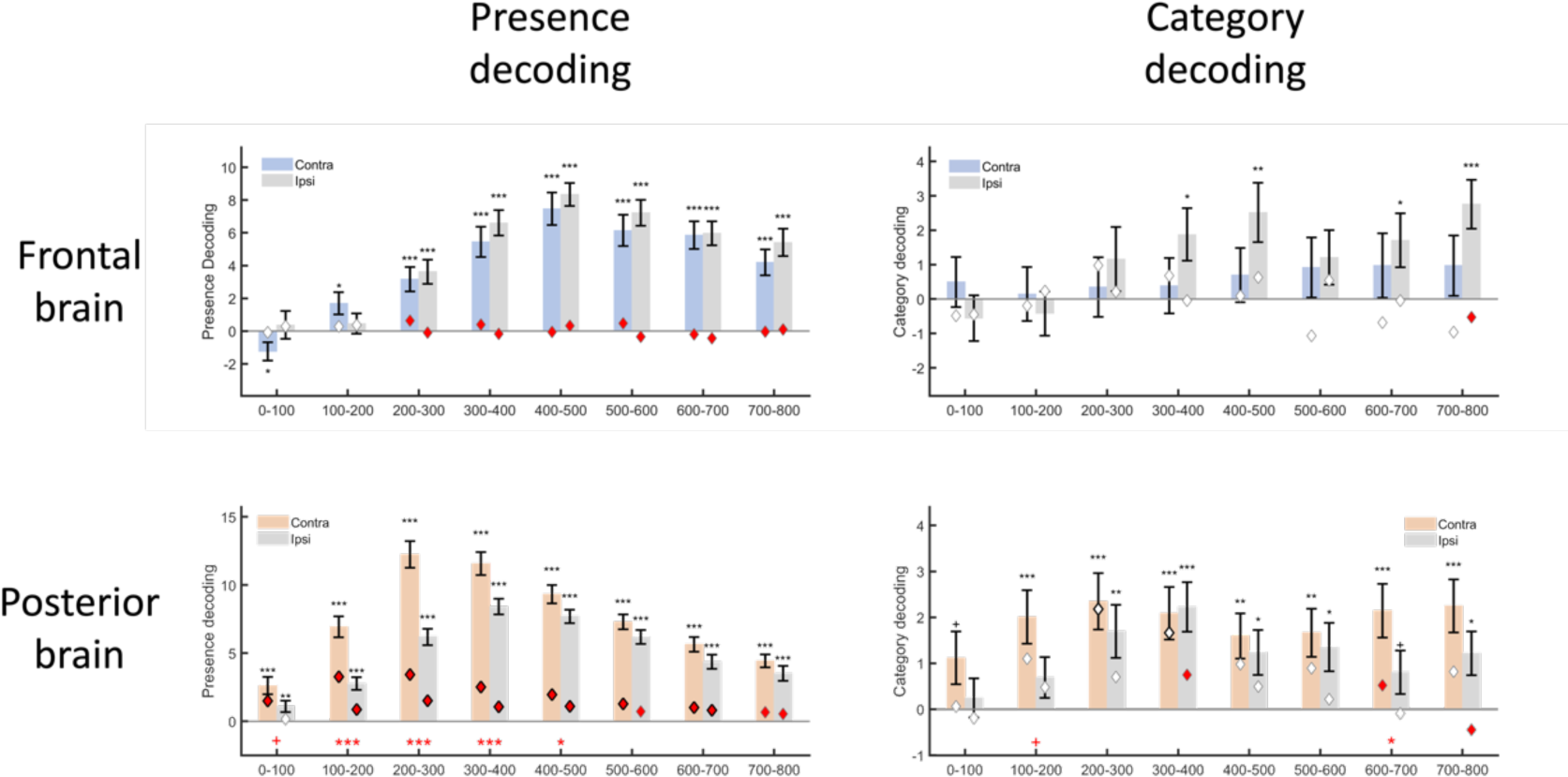
The latency and degree of neural lateralization of the frontal brain (upper row) and the posterior brain (lower row) in decoding coarse-grained (left column) and fine-grained (right column) signals of consciousness. Visibility-related (visible-invisible) decoding performance was averaged across each bin of 100ms for contralateral versus ipsilateral stimuli, respectively. For both types of decoding analysis, features imported into the SVM model were multivariate, including mean ERP response and mean power of six frequency bands ranging from slow delta to fast high gamma oscillations (see Methods for more information). For each panel figure, decoding performance for contralateral versus ipsilateral stimulus was compared, with red asterisk shown at the bottom indicating statistical difference between these two bars in the attended condition. To evaluate decoding performance of each bar, bar values were also compared against 0. For the attended condition, statistical significance of this test was indicated by an asterisk shown above the error bar. For the unattended condition, mean decoding performance was indicated by the position of the diamond symbol and significance of paired t-test against 0 was coded by its edge color. Diamond with bold and black edges indicates above-chance decoding performance (uncorrected p<0.05) in the unattended condition (note that same responsive contacts were used) while the one outlined by thin and gray edges indicates a non-significant result (uncorrected p>0.05). Beyond that, face color of the diamond indicates the statistical difference between attended and unattended condition on decoding performance, with red face color indicating a significant attentional modulation effect (two-tailed, uncorrected p <0.05) while the white face color indicating a non-significant modulation effect (p>0.05). To better evaluate any trend effect, correction for multiple comparisons were not conducted in this figure. +, p<0.1; *, p<0.05; **, p<0.01; ***, p<0.001; otherwise, p>0.1. Error bar indicates SEM across responsive units. Key points of this figure were summarized below. In terms of response latency in decoding the consciousness signal in the attended condition, the posterior brain became responsive within the first 100ms, and temporally led the frontal brain in decoding both coarse-grained representation of consciousness presence and fine-grained representation of consciousness contents. In terms of the degree of neural lateralization, this lateralization effect was pronounced in representing the coarse-grained signal of consciousness presence, rather than fine-grained signal of consciousness contents. Specifically, contralateral hemisphere of the posterior brain conveyed stronger signal than the ipsilateral hemisphere, an effect continuously lasting within the first 500ms. On the contrary, the two hemispheres of the frontal brain operated equally during this process.

**Fig.S18.**
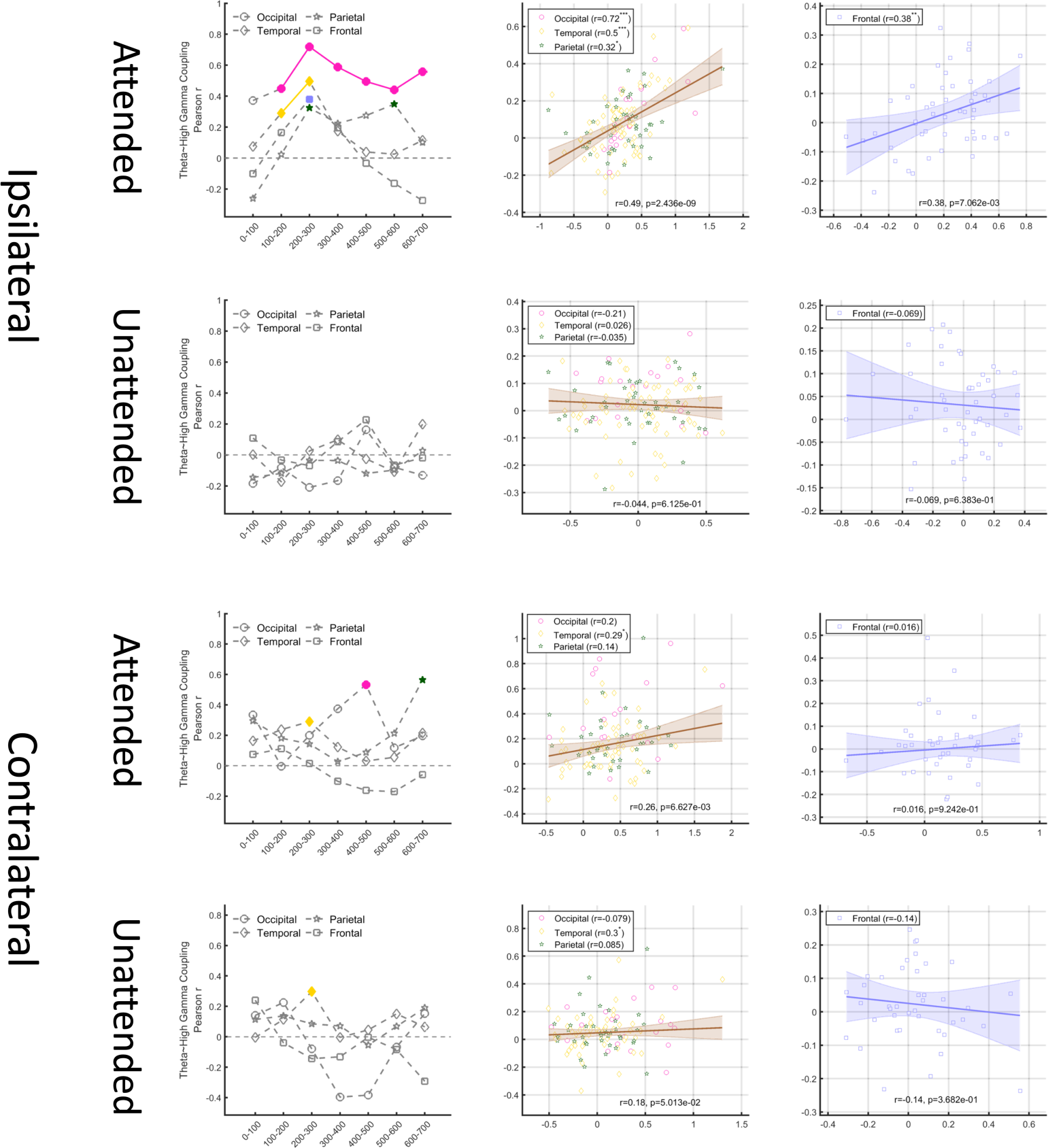
Theta-high gamma cross frequency coupling. As well known, the imported visual information reaches the contralateral posterior hemisphere first. So, the ‘evoked’ high gamma activity at the contralateral hemisphere may largely derive from stimulus-evoked response. By contrast, without such directly evoked response, in the ipsilateral posterior hemisphere high gamma activity may be more intrinsically prone to the consequence of long-distance global coordination via mechanisms like cross-frequency coupling. Based on this knowledge, we hypothesized that if conscious signals are exchanged across the whole brain to coordinate distant local high gamma activity, this mechanism is more likely to be detected in the ipsilateral hemisphere whose high gamma activity is less affected by stimulus-evoked response relative to the contralateral hemisphere. To this end, we calculated theta-high gamma cross frequency coupling (i.e., Pearson correlation between visibility-related measures for theta power and high gamma power) for both the ipsilateral hemisphere (1 and 2 rows) and the contralateral hemisphere (3 and 4 rows) in both attended (odd numbered rows) and unattended (even numbered rows) conditions. The first column shows how this correlation varied across bins of 100ms, with colored symbols indicate significant correlation (FDR corrected p<0.05). The second and third columns show scatterplot for the posterior brain and the frontal brain, respectively, to illustrate the underlying data distribution at the 200-300ms bin where the targeted coupling was mostly pronounced. Based on considerations above, we paid special interest to examine whether theta-high gamma cross frequency coupling occurred in the ipsilateral hemisphere. Expectedly, significant coupling was preferentially observed at the ipsilateral hemisphere, particularly for the occipital and parietal lobes. Also importantly, this correlation was mainly observed in the attended condition, suggesting that attention has the privilege to engage theta oscillations to exchange coarse-grained signals globally to coordinate distant local activities.

**Table S1.**
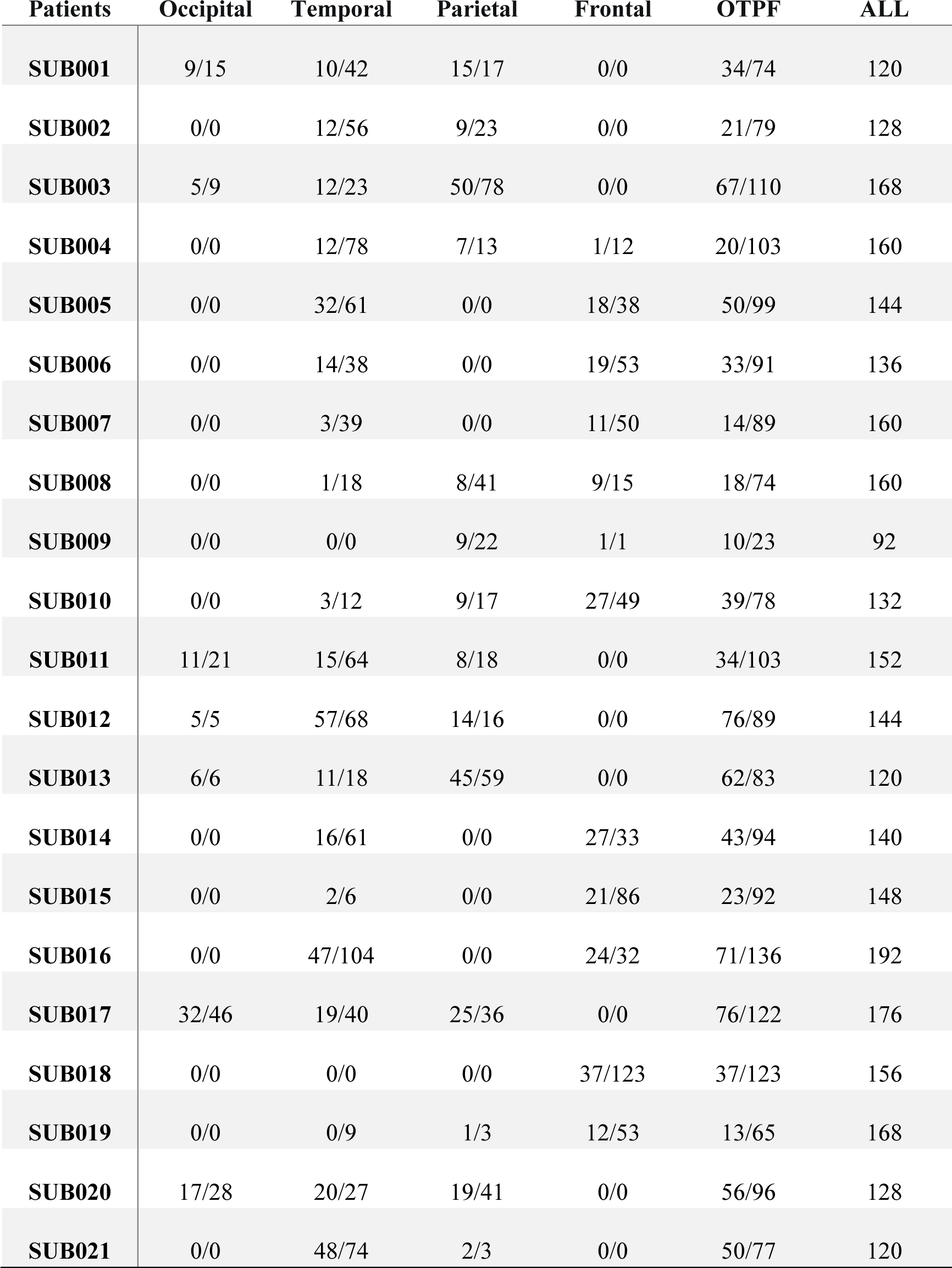
The information regarding the number of responsive contacts.

**Table S2.** The location of responsive contacts at the frontal lobe (n=16) and parietal lobe (n=62) in the unattended condition.

| **ID** | **Lobe** | **Patient ID** | **Electrode ID** | **Contact ID** | **MNIs** |  |  | **Yeo7_label** |
| --- | --- | --- | --- | --- | --- | --- | --- | --- |
| **1** | Frontal | 5 | 5-10 | 690 | -37 | 41 | 10 | Frontoparietal |
| **2** | Frontal | 5 | 5-10 | 691 | -36 | 45 | 13 | Frontoparietal |
| **3** | Frontal | 5 | 5-11 | 705 | -54 | 20 | 15 | DefaultNetwork |
| **4** | Frontal | 6 | 6-04 | 773 | -45 | 33 | 25 | Frontoparietal |
| **5** | Frontal | 7 | 7-06 | 930 | -34 | 37 | 39 | Ventral attention |
| **6** | Frontal | 10 | 10-01 | 1280 | 43 | 33 | -3 | Ventral attention |
| **7** | Frontal | 14 | 14-01 | 1828 | 40 | 50 | -1 | Frontoparietal |
| **8** | Frontal | 14 | 14-02 | 1840 | 45 | 29 | 2 | Ventral attention |
| **9** | Frontal | 14 | 14-02 | 1841 | 46 | 26 | 2 | Ventral attention |
| **10** | Frontal | 14 | 14-02 | 1843 | 50 | 26 | 5 | DefaultNetwork |
| **11** | Frontal | 14 | 14-09 | 1925 | 44 | 27 | 14 | Frontoparietal |
| **12** | Frontal | 15 | 15-02 | 1977 | -29 | 32 | -14 | Limbic |
| **13** | Frontal | 15 | 15-02 | 1978 | -32 | 31 | -11 | DefaultNetwork |
| **14** | Frontal | 15 | 15-03 | 1989 | -20 | 48 | -6 | Frontoparietal |
| **15** | Frontal | 15 | 15-11 | 2099 | 36 | 44 | -5 | Frontoparietal |
| **16** | Frontal | 18 | 18-12 | 2624 | 21 | 53 | 2 | Frontoparietal |
| **17** | Parietal | 2 | 2-02 | 137 | -43 | -51 | 31 | DefaultNetwork |
| **18** | Parietal | 3 | 3-02 | 267 | 19 | -42 | 42 | Ventral attention |
| **19** | Parietal | 3 | 3-04 | 300 | 25 | -63 | 34 | Dorsal attention |
| **20** | Parietal | 3 | 3-04 | 302 | 29 | -66 | 33 | Dorsal attention |
| **21** | Parietal | 3 | 3-05 | 313 | 14 | -58 | 23 | DefaultNetwork |
| **22** | Parietal | 3 | 3-05 | 316 | 23 | -63 | 27 | Visual |
| **23** | Parietal | 3 | 3-05 | 318 | 31 | -65 | 25 | Dorsal attention |
| **24** | Parietal | 4 | 4-11 | 556 | -30 | -50 | 50 | Dorsal attention |
| **25** | Parietal | 8 | 8-03 | 1049 | 17 | -40 | 60 | Somatomotor |
| **26** | Parietal | 8 | 8-12 | 1166 | 22 | -52 | 13 | Visual |
| **27** | Parietal | 8 | 8-12 | 1174 | 51 | -56 | 29 | DefaultNetwork |
| **28** | Parietal | 10 | 10-08 | 1361 | 31 | -29 | 17 | Somatomotor |
| **29** | Parietal | 11 | 11-09 | 1517 | 39 | -57 | 18 | DefaultNetwork |
| **30** | Parietal | 11 | 11-09 | 1518 | 42 | -57 | 26 | DefaultNetwork |
| **31** | Parietal | 11 | 11-09 | 1519 | 44 | -56 | 25 | DefaultNetwork |
| **32** | Parietal | 12 | 12-07 | 1633 | 8 | -51 | 13 | DefaultNetwork |
| **33** | Parietal | 12 | 12-07 | 1635 | 19 | -55 | 10 | Visual |
| **34** | Parietal | 12 | 12-08 | 1656 | 43 | -63 | 20 | Dorsal attention |
| **35** | Parietal | 12 | 12-08 | 1657 | 42 | -64 | 19 | Dorsal attention |
| **36** | Parietal | 13 | 13-01 | 1702 | -17 | -46 | 39 | DefaultNetwork |
| **37** | Parietal | 13 | 13-01 | 1710 | -35 | -54 | 54 | Dorsal attention |
| **38** | Parietal | 13 | 13-01 | 1711 | -38 | -53 | 55 | Dorsal attention |
| **39** | Parietal | 13 | 13-02 | 1720 | -26 | -69 | 28 | Dorsal attention |
| **40** | Parietal | 13 | 13-02 | 1721 | -29 | -71 | 29 | Dorsal attention |
| **41** | Parietal | 13 | 13-02 | 1722 | -34 | -71 | 35 | Dorsal attention |
| **42** | Parietal | 13 | 13-02 | 1723 | -37 | -75 | 39 | DefaultNetwork |
| **43** | Parietal | 13 | 13-02 | 1724 | -38 | -77 | 40 | DefaultNetwork |
| **44** | Parietal | 13 | 13-02 | 1725 | -39 | -75 | 42 | DefaultNetwork |
| **45** | Parietal | 13 | 13-02 | 1726 | -41 | -75 | 42 | DefaultNetwork |
| **46** | Parietal | 13 | 13-02 | 1727 | -43 | -74 | 46 | DefaultNetwork |
| **47** | Parietal | 13 | 13-03 | 1737 | -27 | -75 | 19 | Visual |
| **48** | Parietal | 13 | 13-03 | 1738 | -30 | -78 | 19 | Visual |
| **49** | Parietal | 13 | 13-03 | 1741 | -39 | -84 | 26 | Dorsal attention |
| 50 | Parietal | 13 | 13-03 | 1742 | -41 | -83 | 27 | Dorsal attention |
| 51 | Parietal | 13 | 13-03 | 1743 | -42 | -83 | 31 | Dorsal attention |
| 52 | Parietal | 13 | 13-04 | 1752 | -25 | -63 | 27 | Dorsal attention |
| 53 | Parietal | 13 | 13-04 | 1753 | -31 | -65 | 28 | Dorsal attention |
| 54 | Parietal | 13 | 13-04 | 1758 | -44 | -67 | 44 | DefaultNetwork |
| 55 | Parietal | 13 | 13-04 | 1759 | -44 | -64 | 49 | DefaultNetwork |
| 56 | Parietal | 13 | 13-04 | 1760 | -48 | -67 | 46 | DefaultNetwork |
| 57 | Parietal | 13 | 13-05 | 1768 | -28 | -81 | 8 | Visual |
| 58 | Parietal | 13 | 13-05 | 1769 | -32 | -81 | 9 | Visual |
| 59 | Parietal | 13 | 13-05 | 1770 | -35 | -82 | 9 | Visual |
| 60 | Parietal | 13 | 13-05 | 1772 | -42 | -81 | 15 | Visual |
| 61 | Parietal | 13 | 13-06 | 1782 | -39 | -63 | 11 | Dorsal attention |
| 62 | Parietal | 13 | 13-06 | 1783 | -41 | -62 | 11 | Dorsal attention |
| 63 | Parietal | 17 | 17-03 | 2341 | 44 | -76 | 19 | Visual |
| 64 | Parietal | 17 | 17-03 | 2342 | 47 | -76 | 22 | Dorsal attention |
| 65 | Parietal | 17 | 17-03 | 2343 | 49 | -73 | 25 | Dorsal attention |
| 66 | Parietal | 17 | 17-04 | 2358 | 46 | -70 | 11 | Visual |
| 67 | Parietal | 17 | 17-10 | 2428 | 12 | -40 | 42 | Ventral attention |
| 68 | Parietal | 17 | 17-11 | 2450 | -31 | -84 | 8 | Visual |
| 69 | Parietal | 17 | 17-11 | 2452 | -38 | -86 | 11 | Visual |
| 70 | Parietal | 17 | 17-12 | 2466 | -35 | -77 | 21 | Visual |
| 71 | Parietal | 17 | 17-12 | 2467 | -36 | -79 | 26 | Visual |
| 72 | Parietal | 17 | 17-12 | 2468 | -35 | -82 | 30 | Dorsal attention |
| 73 | Parietal | 17 | 17-12 | 2469 | -35 | -83 | 34 | Dorsal attention |
| 74 | Parietal | 20 | 20-01 | 2805 | 37 | -41 | 43 | Dorsal attention |
| 75 | Parietal | 20 | 20-02 | 2824 | 39 | -71 | 49 | Frontoparietal |
| **76** | Parietal | 20 | 20-03 | 2831 | 27 | -72 | 26 | Visual |
| **77** | Parietal | 20 | 20-04 | 2845 | 11 | -77 | 29 | Visual |
| **78** | Parietal | 20 | 20-05 | 2855 | 23 | -77 | 16 | Visual |

For contacts of the same electrode ID, their spatial neighborhood could be inferred by Contact ID. For example, contact 2468 and 2469 were two neighboring contacts from the same electrode (electrode ID: 17-12, i.e., electrode 12 of patient 17) while contact 2452 was from a different electrode of the same patient (electrode ID: 17-11).

**Movie S1.**

Demo illustration of trials without any probes displayed in the pretest session (number-counting task) of Exp.1.

**Movie S2.**

Demo illustration of trials with randomly selected occasional probes displayed in the posttest session (number-counting task) of Exp.1.

**Movie S3.**

Demo illustration of trials displayed in the unattended blocks (number-counting task) of Exp.2.

**Movie S4.**

Demo illustration of trials displayed in the attended blocks (image-categorization task) of Exp.2.

**Movie S5.**

Spatial distribution of responsive contacts across four neocortex lobes and across time.

